# Epigenetic delineation of the earliest cardiac lineage segregation by single-cell multi-omics

**DOI:** 10.1101/2024.05.17.594655

**Authors:** Peng Xie, Xu Jiang, Jingjing He, Qingyun Pan, Xianfa Yang, Yanying Zheng, Zhuanzhuan Che, Wenli Fan, Chen Wu, Weiheng Zheng, Shuhan Si, Kun Gao, Shiqi Zhu, Ke Fang, Haitong Fang, Yi Yang, Tao P. Zhong, Zhongzhou Yang, Ke Wei, Wei Xie, Naihe Jing, Zhuojuan Luo, Chengqi Lin

## Abstract

The mammalian heart is formed from multiple mesoderm-derived cell lineages. However, it remains largely unknown when and how the specification of mesoderm towards cardiac lineages is determined. Here, we systematically depict the transcriptional trajectories toward cardiomyocyte in early mouse embryo, and characterize the epigenetic landscapes underlying the early mesodermal lineage specification by single-cell multi-omics analyses. The analyses also reveal distinct core regulatory networks (CRN) in controlling specification of mesodermal lineages. We further demonstrate the essential role HAND1 and FOXF1 in driving the earliest cardiac progenitors specification. These key transcription factors occupy at distinct enhancers, but function synergistically and hierarchically to regulate the expression of cardiac-specific genes. In addition, HAND1 is required for exiting from the nascent mesoderm program, while FOXF1 is essential for driving cardiac differentiation during juxta-cardiac field (JCF) specification. Our findings establish transcriptional and epigenetic determinants specifying the early cardiac lineage, providing insights for the investigation of congenital heart defects.

## Introduction

Heart development requires coordinated specification of multiple lineages, each characterized by serial cell fate determination events. Identifying the developmental trajectories is the key to understanding heart formation [1, 2]. In past decades, a stepwise determination of early cardiac lineage hierarchy has been primarily established [3]. The current model designates cardiac progenitor cells into discrete pools, including the first and second heart field (FHF and SHF) [1], and a newly classified juxta-cardiac field (JCF) [4]. FHF mainly contributes to the left ventricle (LV) and the atria. SHF, located in a dorsal-medial region to FHF, progressively develops into cells in the right ventricle (RV), the outflow tract (OFT) and the atria [5]. Interestingly, JCF contributes to not only epicardium and pericardium, but also cardiomyocytes (CMs) of LV and the atria [4, 6].

Cardiac progenitors are mostly generated from the *Mesp1*-expressing (*Mesp1^+^*) nascent mesoderm (NM) cells during gastrulation [7, 8]. Pioneering work has revealed that the developmental capacities of each cardiac progenitor pool are highly related to the spatial-temporal constriction during the specification of NM cells [6, 9, 10]. Temporally inducible lineage tracing indicates that E6.5 *Mesp1^+^* cells mostly contribute to LV, whereas E7.25 *Mesp1^+^* cells give rise to RV, atria, OFT, and inflow tracts (IFT) [9]. It seems that early-streak stage NM cells differentiate into FHF pools, while late-streak stage NM cells relate to SHF progenitors. Interestingly, JCF population is also derived from the *Mesp1^+^*NM cells in the gastrula [6]. Recent studies on single-cell transcriptomic data of the late headfold stage embryos have revealed that JCF shares a number of molecular markers with FHF, but lacks *Nkx2.5* expression and exhibits specific *Mab21l2* expression [4]. However, unlike FHF cells, JCF cells are largely located at the embryonic-extraembryonic mesodermal interface, as revealed by the *Mab21l2* expression, rostrally to the *Nkx2-5* positive cardiac crescent region [4]. It remains unclear the molecular signaling underlying the early specification of NM cells into JCF and FHF population.

In this study, by bridging the transcriptional landscapes between the gastrula and the headfold stage in early mouse embryos, we systematically depict the transcriptional trajectories leading to CMs during early mouse development, and characterize the epigenetic landscapes that underlie early mesodermal lineage specification. The analyses reveal two dinstinct developmental trajectories towards CMs, namely the JCF trajectory (the *Hand1*-expressing early extraembryonic mesoderm – JCF and FHF – CM) and the SHF trajectory (the pharyngeal mesoderm cells – SHF – CM). Through single-cell multi-omics analysis, a predicted core regulatory network (CRN) in JCF is identified, consisting of transcription factors (TFs) GATA4, TEAD4, HAND1 and FOXF1. Further functional analysis indicates that HAND1 and FOXF1 are activated sequentially, and both required for mesodermal specification and the expression of the JCF specific genes. Taken together, our study unveils the transcriptional and epigenetic dynamics during early cardiac specification, demonstrates the crucial roles of HAND1 and FOXF1 in driving early cardiac specification, and provides insights for the investigation of congenital heart defects.

## Results

### Transcriptional dynamics during the specification of early cardiac progenitors

To delineate the origin of cardiac progenitors, we constructed the E6.5-8.5 developmental trajectories using the published mouse gastrulation cell atlas by performing the Waddington-Optimal-Transport (WOT) analysis to infer ancestor-descendant fates of cells [11, 12]. WOT models the temporal dynamics of cell state transitions by computing probabilistic couplings between adjacent timepoints, effectively inferring ancestor-descendant relationships based on transcriptomic similarity and temporal continuity. Here, CMs were used as trajectory endpoints and traced back to the E6.5 epiblast. Clear trajectory separation was observed within E7.5-7.75 (Figure 1A). Besides the pharyngeal mesoderm (PM) cell cluster at E7.75, a subset of E7.5 early extraembryonic mesoderm (EEM) cells [6] was specifically identified in a distinct developmental trajectory. PM population was marked by the expression of *Isl1*, *Sfrp1*, *Tcf21*, *Tbx1* and *Irx3*, suggesting its relationship with the SHF progenitors; the EEM cells exhibited highly expressing *Hand1*, *Pmp22*, *Foxf1* and *Spin2c* (Figure S1A). In addition, cells along the EEM trajectory also expressed higher levels of the FHF signature genes, including *Tbx5* and *Hcn4*, suggesting their contribution to the FHF [6]. Compared with the PM trajectory, the EEM trajectory was mainly composed of cells in the later stages of cardiac development (Figure 1B and S1B). By the heart looping stage (E8.5), the two trajectories indeed exhibited distinct contributions to cardiac structures LV and OFT, respectively (Figure 1C).

**Figure 1:**
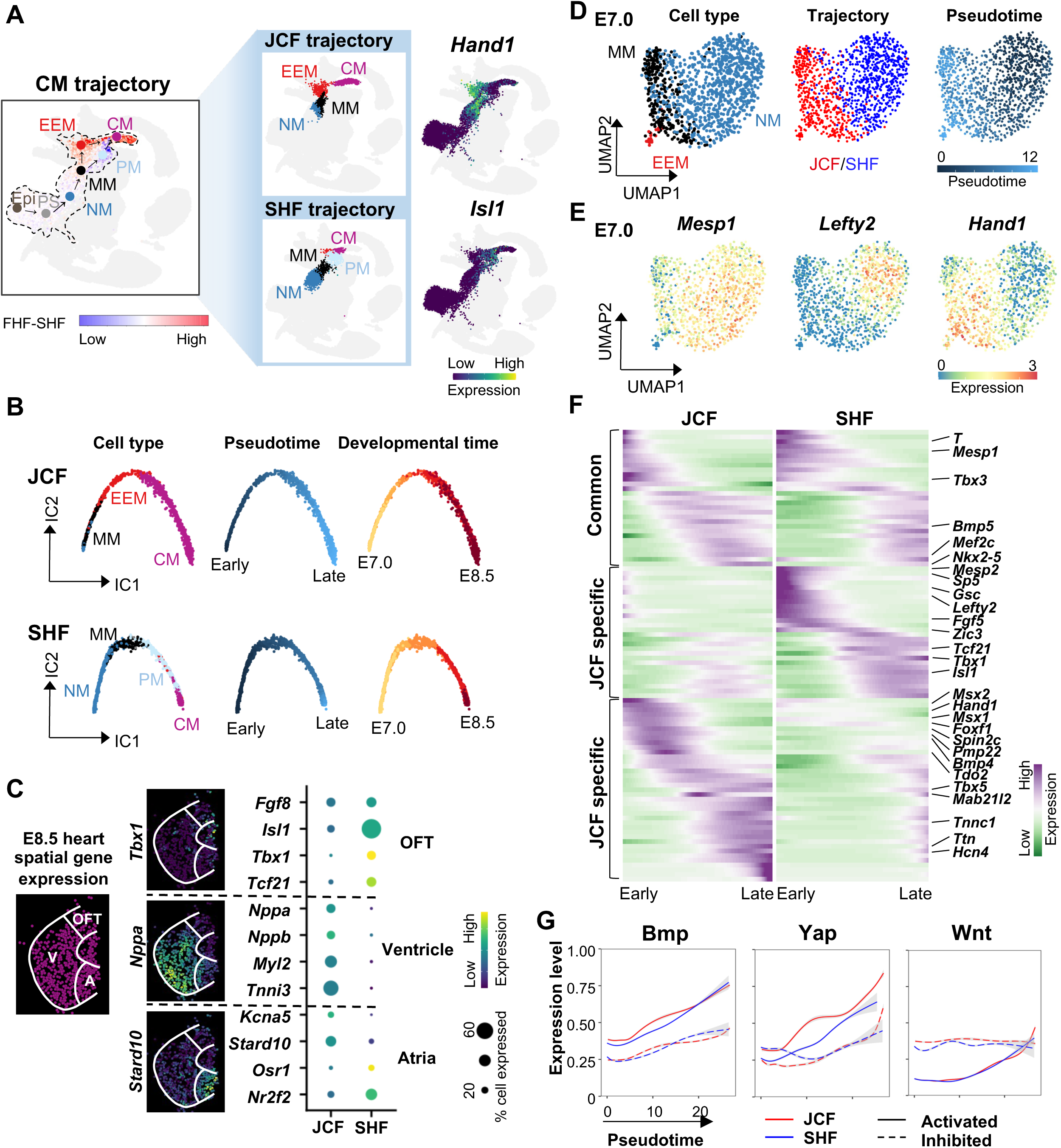
Inferred trajectories reflect two distinct development routes of CMs. **(A)** Two distinct trajectories, the JCF trajectory and the SHF trajectory, were inferred during CM differentiation. UMAP layout from Pijuan-Sala et al. [12] is highlighted by cells belonging to the WOT predicted developmental trajectories for CM (left), EEM (upper middle) and PM (lower middle), respectively. UMAP layout for the CM trajectory with cells is colored by the difference between FHF- and SHF-gene signature scores. UMAP layout for the JCF or SHF trajectory is colored by cell types. Gene expression levels of *Hand1* (upper right) and *Isl1* (lower right) of cells along the CM trajectory is overlaid onto the above-mentioned UMAP layout. Epi, epiblast. PS, primitive streak. NM, nascent mesoderm. MM, mixed mesoderm. EEM, early extraembryonic mesoderm. PM, pharyngeal mesoderm. CM, cardiomyocyte. **(B)** Independent component (IC) layout showing pseudotemporal trajectories for JCF trajectory (upper) and SHF trajectory (lower) cells, colored by cell type (left), pseudotime (middle) and developmental time (right). **(C)** The JCF and SHF trajectories showing distinct contributions to cardiac structures. Spatial plot showing spots in cardiac subregions from E8.5 embryos (left) [29]. White lines denoting the region including outflow tract (OFT), ventricle (V) and Atria (A). ‘Virtual’ in situ hybridization (vISH) confirming spatial specificity of marker genes corresponding to cardiac subregions (middle). Dot plot showing expression difference in subregion specific genes in the JCF and SHF trajectories (right). **(D)** Mesodermal lineage segregation at E7.0. UMAP layout for E7.0 CM trajectory cells is colored by cell type (left), trajectory (middle) and pseudotime (right). **(E)** UMAP layout (same as d) showing the expression of the E7.0 marker genes of NM (*Mesp1*), JCF (*Hand1*) and SHF (*Lefty2*). **(F)** Heatmap showing the expression of pseudotime-dependent genes for the JCF trajectory (right) and the SHF trajectory (left). Rows and columns represent genes and cell bins, respectively. Genes used for the heatmap are listed in Supplementary Table 1. **(G)** Smoothened fitting curves showing expression levels of activated (solid line) and inhibited (dotted line) signaling markers in the JCF (red) and SHF (blue) trajectories.

Thus, we here refer to these two transcriptional trajectories as JCF and SHF (Figure 1A). The spatial correlation of these two trajectories with heart fields was confirmed using the dataset from manually micro-dissected mesodermal cells in cardiac regions of E7.75-8.25 mouse embryos (Figure S1C) [4]. The JCF and SHF trajectories contained overlapping but temporally distinct cell types (Figure 1B and Figure S1D). We performed analysis of fate divergence between two trajectories, which suggests, before E7.0, mesodermal cells have similar probabilities to choose either trajectory (Figure S1E).The separation of the two trajectories can be observed as early as E7.0 (Figure 1B and Figure S1E). NM cells, marked by *Mesp1* and *Lefty2*, at E7.0 were more likely to be the multipotent progenitors of the SHF trajectory; whereas mixed mesoderm (MM) cells, marked by *Hand1* and *Msx2*, in later developmental stage and EEM cells tended to contribute to the JCF trajectory (Figures 1D, 1E and Figure S1F). To illucidate the contribution of JCF and SHF lineages to the cardiac crescent (CC), we used early CC progenitor population (E7.75 *Nkx2- 5*+; *Mab21l2*- CMs) as the starting point and performed WOT lineage inference (Figure S2A). Results suggest that both JCF and SHF progenitors contribute to the CC, consistent with live imaging-based single cell tracing by Dominguez et al [13] and lineage tracing results by Zhang et al [6]. We also analyzed the expression levels of CC marker genes (*Tbx5*, *Hcn4*) and observed their activation along both trajectories (Figure S2B).

We then systematically characterized the stage specific marker genes in the JCF and SHF trajectories. These two trajectories exhibited discrete and dynamic gene expression profiles during development (Figure 1F). The JCF marker gene, *Mab21l2*, showed transient expression [4], in contrast to the late but continued expression of SHF markers (*Isl1*, *Tbx1*), further supporting the asynchronized fate commitment by the two lineages. Notably, along the pseudotime trajectory, the NM marker *Mesp1* displays a transient and early expression in the JCF lineage, whereas it is persistently expressed during the mid-to-late stages in the SHF lineage, which suggests distinct mechanisms of mesodermal specification underlying the two lineages. We also observed relatively similar inhibitory Wnt and Nodal, as well as active Fgf and Notch, signaling activities along the two trajectories (Figures 1G and S3A). Interestingly, the early stage of the JCF trajectory seems show higher Bmp and Yap signaling activities (Figure 1G). Temporal expression profiles of the *Bmp* genes indicated that *Bmp4*/*5*/*7* were dynamically expressed during cardiac specification, with *Bmp4* demonstrating higher JCF specificity and at least 0.5 days earlier activation (Figure S3B). Geo-seq data analysis indicated that *Bmp4* was highly specific to mesoderm, and enriched at the proximal mesodermal ends (layer 11 at E7.0, layer 9-10 at E7.5) with distinct anterior-posterior preference at E7.5 (Figure S3C). For the target genes of Bmp signaling, several genes (*Hand1*, *Car4*, *Arl4c* and *Pmp22*) showed JCF specific activation-to-repression dynamics, similar to *Bmp4* (Figures 1F and S3D).

### Epigenetic signatures of the early JCF and SHF progenitors

To investigate the epigenetic regulation during cardiac cell fate decisions, we performed multi-omic analysis by combining single-nucleus RNA-sequencing (snRNA-seq) and scATAC-seq to generate paired, cell type specific transcriptome and chromatin accessibility profiles of 13,226 cells in E7.0 mouse embryos. The single-cell transcriptomic data were integrated with the published E7.0 mouse embryo cell atlas data, followed by label transfer and gene expression-based cell type identification (Figures S4A and S4B). For the scATAC-seq data, we scored the genome-wide ATAC activities with bin sizes of 10 kb prior to UMAP analysis, which yielded cell clusters similar to transcriptome-based analysis (Figure S4A).

Nine clusters of cells were identified through clustering analysis at both transcriptional and epigenetic levels, which are NM cells (Clusters 0, 1 and 2), primordia germ cells (PGC, Cluster 4), hematoendothelial progenitors (Haem, Clusters 5 and 6), and EEM cells (Clusters 3, 7 and 8) (Figures 2A, 2B and S4C). RNA velocity also supported the four possible trajectories mentioned above for the earliest mesodermal cell specification (Figure 2C). WOT analysis revealed that Clusters 3, 7, and 8 showed intermediate to high probabilities of belonging to the JCF trajectory; pseudotime analysis indicated that Cluster 8 represented the late differentiated EEM populations (Figure S5A). Although Cluster 2 represented the relatively late stage of NM cell population by pseudotime analysis, Cluster 1, 0 and 2 demonstrated similar probabilities of belonging to the SHF trajectory (Figure S5A). Thus, the analyses further indicated that EEM cells were clearly separated from the nascent mesoderm, while the SHF trajectory related cells still remained at the early NM stage at E7.0.

**Figure 2:**
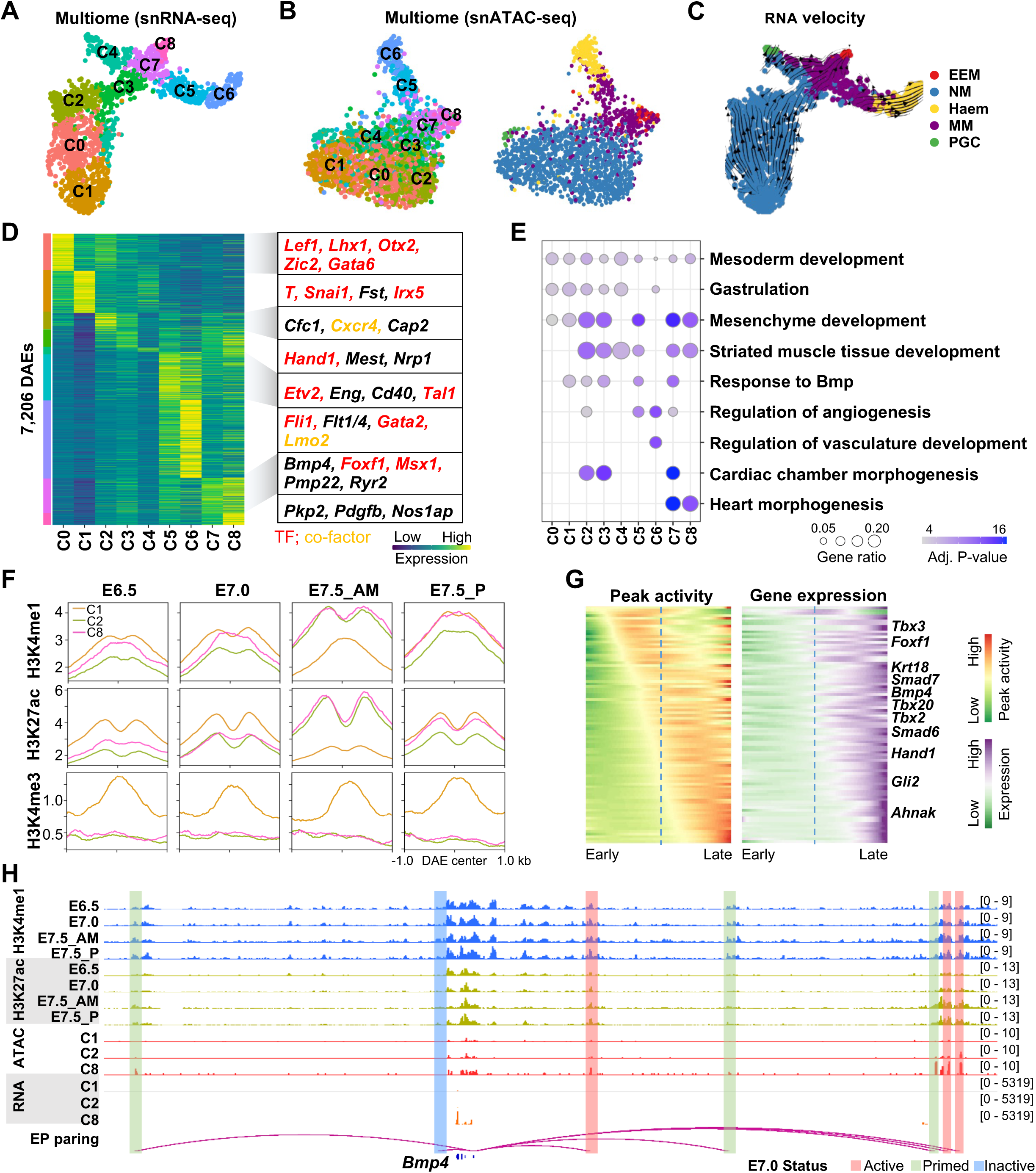
Multi-omics analysis reveals epigenetic signatures of the early JCF and SHF progenitors. **(A-C**) Clustering analysis of E7.0 single-cell multi-omics data and comparison between modalities of transcriptome (snRNA-seq) and chromatin accessibility (snATAC-seq). UMAP layout, using only snRNA-seq (A,C) or snATAC-seq (B) data, is colored by cluster identities (left) or cell types (right). For both (A) and (B), cluster identities are determined by snRNA-seq data. For (C), developmental directions, shown as arrows, are indicated by RNA-velocity analysis. **(D)** Heatmap showing the activity of 7,206 differentially accessible elements (DAEs) between clusters. For the DAE motif analysis, we firstly inferred the motif and TF families, then tested which specific TFs are expressed in the corresponding cell cluster. Rows and columns represent DAEs and clusters. Colors indicate levels of accessibility averaged among cells from each cluster. The length of the corresponding color bar on the left represents the number of DEAs. Representative DAE-associated genes are shown on the right side. **(E)** Dot plot indicating functional enrichment of DAE-associated genes of each cluster. **(F)** Epigenetic status of DAEs during mesodermal specification of E6.5-7.5 mice embryo. H3K4me1, H3K27ac and H3K4me3 profiles were averaged among DAEs of cluster C1 (NM), C2 (SHF-direction frontier) and C8 (JCF-direction frontier). Published H3K4me1, H3K27ac and H3K4me3 ChIP-seq data were collected from E6.5 epiblast, E7.0/7.5 posterior epiblast (P) and E7.5 anterior mesoderm (AM) [14]. **(G)** Smoothened heatmap showing dynamic gene expression (right) and enhancer accessibility (left) along E7.0 mesoderm pseudotime trajectories for gene-enhancer pairs. Dashed lines indicating the pseudo-temporal midpoint. **(H)** Representative genome browser snapshots of ATAC/RNA-seq (aggregated gene expression and chromatin accessibility for each cluster), H3K4me1 and H3K27ac at the *Bmp4* locus. Putative enhancers status at E7.0 are highlighted by colors.

We further analyzed the chromatin accessibilities of these cell clusters. Total 90,661 chromatin accessible elements (CAEs) were detected, 7,206 of which were differentially accessible elements (DAEs) across the 9 clusters (Figure 2D). The DAEs were annotated to their target genes by enhancer-promoter (EP) pairing analysis. Consistent with the clustering analysis based on gene expression (Figure S4C), DAVID functional term analysis revealed that the DAE target genes in Clusters 3, 7 and 8 , such as *Hand1*, *Foxf1*, *Bmp4* and *Msx1*, were mainly associated with heart morphogenesis, that those in Clusters 5 and 6, like *Tal1*, *Lmo2* and *Fli1*, were related to angiogenesis and vasculature development, and that those in Clusters 0, 1, 2, for examples, *T*, *Zic2*, *Lhx1* and *Gata6*, were enriched in gastrulation and mesoderm development (Figures 2D and 2E).

To characterize the spatiotemporal chromatin dynamics of the DAEs in JCF and SHF, we quantified the occupancies of the enhancer marks H3K4me1 and H3K27ac, as well as the promoter mark H3K4me3 at these DAEs across the E6.5-7.5 developmental stages[14]. JCF/Cluster 8 and SHF/Cluster 2 specific DAEs could potentially function as enhancers and become activated at anterior regions of E7.5 embryos, as the DAEs were generally marked by H3K27ac and H3K4me1, but not H3K4me3 (Figure 2F). The enrichment of H3K4me1 at E7.0 and even earlier at E6.5 stage, along with the higher levels of H3K27ac at these DAEs at E7.5 stage, suggested that many of these DAEs could be dormant or inactive enhancers during earlier stages like E6.5, but primed for later activation during lineage specification (Figure 2F). Indeed, the integrated analysis on chromantin accessibility of the DAEs, shown by ATAC-seq, and their target gene expression levels supported that a large portion of the JCF/Cluster 8 DAEs were primed before the full activation of their target genes, like *Bmp4*, *Hand1* and *Foxf1* (Figure 2G). For example, seven DAEs associated with the *Bmp4* gene were identifited, three of which were primed at E7.0 as marked by low levels of H3K27ac but high levels of H3K4me1 (Figure 2H). Taken together, the combined transcriptome and chromatin accessability analysis further supported the early lineage segregation of JCF and the epigenetic priming at gastrulation stage for early cardiac genes.

### Identification of lineage specific key TFs

An integrated analysis of motif enrichment at the DAEs and TF expression data allowed us to identify potential lineage specific key TFs. The SHF/Cluster 2 specific DAEs showed motif enrichment similar to the recognition sequences of known NM specific TFs, including GATA4, ZIC3, EOMES, OTX2, and LHX1 (Figure S5B). Binding sites for hematoendothelium-related TFs, such as GATA2, FLI1, JUNB, and SOX7, were enriched in the hematoendothelial progenitors/Cluster 6 DAEs. In contrast, the binding motifs of GATA4, HAND1, FOXF1 and TEAD4 were highly over-represented in the JCF/Cluster 8 specific DAEs (Figure 3A). Among those JCF-related TFs, GATA4 and TEAD4 showed similar expression and motif activities at Clusters 2, 7, 8 of both JCF and SHF lineages. HAND1 and FOXF1 demonstrated both strong motif activities and specific expression at Cluster 7 and Cluster 8. Interestingly, the expression of HAND1 and FOXF1 seemed relatively transient at NM and EEM cells, and then became downregulated at CM of E7.75 (Figure S5C).

**Figure 3:**
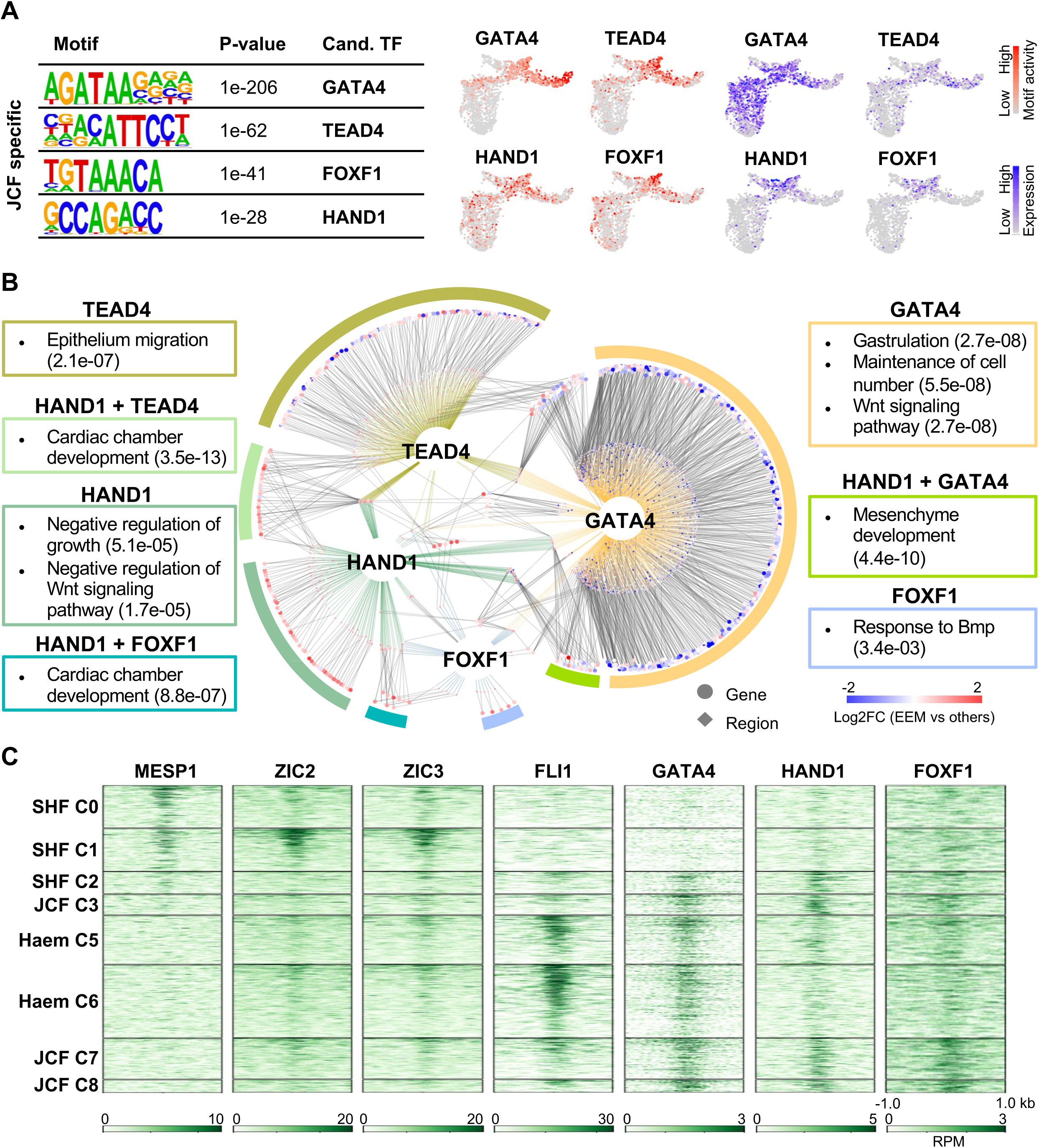
The CRN is identified centering on GATA4, HAND1, FOXF1 and TEAD4 in driving JCF specification. **(A)** Identification of top JCF specific DNA-binding motifs and corresponding candidate TFs. The JCF specific DAEs were defined by comparing C8 with C1 snATAC-seq data using SnapATAC [30] ‘findDAR’ function. Motif calling was performed by the HOMER [31] ‘findMotifsGenome.pl’ function. Motif activity (colored in red) and TF expression (colored in blue) levels of trajectory specific candidate TFs, are overlaid on the UMAP layout from Figure 2A. **(B)** Core regulatory network (CRN) of E7.0 EEM cells. The network is composed of TFs (GATA4, HAND1, FOXF1 and TEAD4), TF binding regions (diamond shapes), target genes (dot shapes), colored lines (TFs to regions) and grey lines (regions to genes). Colors of genes and regions representing the log2 FCs of E7.0 EEM cells over other mesodermal cells, measured by scRNA-seq and snATAC-seq data, respectively. Functional enrichment terms of target genes are shown in boxes with subtitles indicating corresponding TFs. *Q*-values, using hyper-geometric tests, are shown in parenthesis. **(C)** Heatmaps representing the enrichment of MESP1, ZIC2, ZIC3, FLI1, GATA4, HAND1 and FOXF1 across the DAEs of each cluster.

Based on E7.0 single-cell multi-omics data analysis, we predicted a core regulatory network (CRN) centering on the four TFs (Figure 3B). Functional enrichment analyses indicated that this CRN could control key aspects of JCF specification, including Wnt signaling, epithelium cell migration, cell number maintenance, mesenchyme development, and cardiac development. In the CRN, GATA4 and TEAD4 controlled larger gene sets related to the transition from epiblast to mesodermal status, necessary for both JCF and SHF. HAND1 and FOXF1 co-regulated functionally more specific gene sets critical for differentiation to EEM status (Figure 3B). Consistently, most HAND1 and FOXF1 target genes were specifically expressed in EEM, in contrast to GATA4 and TEAD4 target genes (Figure 3B).

We also performed Chromatin immunoprecipitation sequencing (ChIP-Seq) analysis to profile the chromatin occupancies of HAND1 and FOXF1 in mesoderm (MES) and cardiac progenitor (CP) cells derived from the step-wise directed cardiomyocyte differentiation of mouse embryonic stem cells[15]. We also collected published GATA4 ChIP-seq data[16] of E12.5 mouse embryonic heart, FLI1, ZIC2, ZIC3 and MESP1 ChIP-seq data[17] of 2.5 days EB differentiation. Direct comparison of ChIP-seq occupancy profiles with DAEs confirmed the specific enrichment of GATA4, HAND1, and FOXF1 at clusters 7 and 8 of JCF lineages, Fli1 at Clusters 5 and 6, while MESP1 is specifically enrichment at mesoderm cell clusters (Figure 3C). We also noticed the enrichment of HAND1 at early Cluster 3-specific DAEs, while FOXF1 tends to show more speicifc enrichment at cardiac specific enhancers at later CP cells.

### HAND1 and FOXF1 regulate the JCF specific genes

In order to further investigate the molecular roles of HAND1 and FOXF1 in JCF specification, we generated *Hand1* and *Foxf1* KO mouse embryonic stem cell (mESC) lines (Figures S6A-D), followed by *in vitro* mesoderm differentiation and RNA-seq analyses. 2,331 down-regulated and 1,714 up-regulated genes in *Hand1* KO mesoderm (MES) cells, and 870 down-regulated and 970 up-regulated genes in *Foxf1* KO MES cells with fold-change (FC) > 1.5 and *P*-value < 1e-5 were identified (Supplementary Table 2). To explore whether HAND1 and FOXF1 are required for the proper expression of the JCF related genes, we examined the expression of the signature genes of the 9 cell clusters in control , *Hand1* KO and *Foxf1* KO MES cells. First of all, over 90% of the cluster specific genes were detected in the control MES transcriptome, indicating that the *in vitro* differentiation model could be a reliable tool to study the regulation of these cluster specific signature genes (Figure 4A). Indeed, whole transcriptome cosine similarity analysis revealed that the *in vitro* differentiated MES cells were more close to MM and EEM cell state transcriptome-wide (Figure 4B). *Hand1* KO and *Foxf1* KO led to down-regulated expression of a large portion of the Clusters 3, 7 and 8 JCF marker genes, but up-regulated expression of many of the Clusters 0 and 1 SHF marker genes in MES cells (Figure 4A). Consistently, the cosine similarity suggested that *Hand1* and *Foxf1* depletion could lead to EEM differentiation defects (Figure 4B). In addition, many of the down-regulated JCF specific genes and up-regulated SHF specific genes were directly bound by HAND1 and FOXF1 (Figures 4A and S6E). The analysis indicated that HAND1 and FOXF1 were able to directly activate the JCF specific genes.

**Figure 4:**
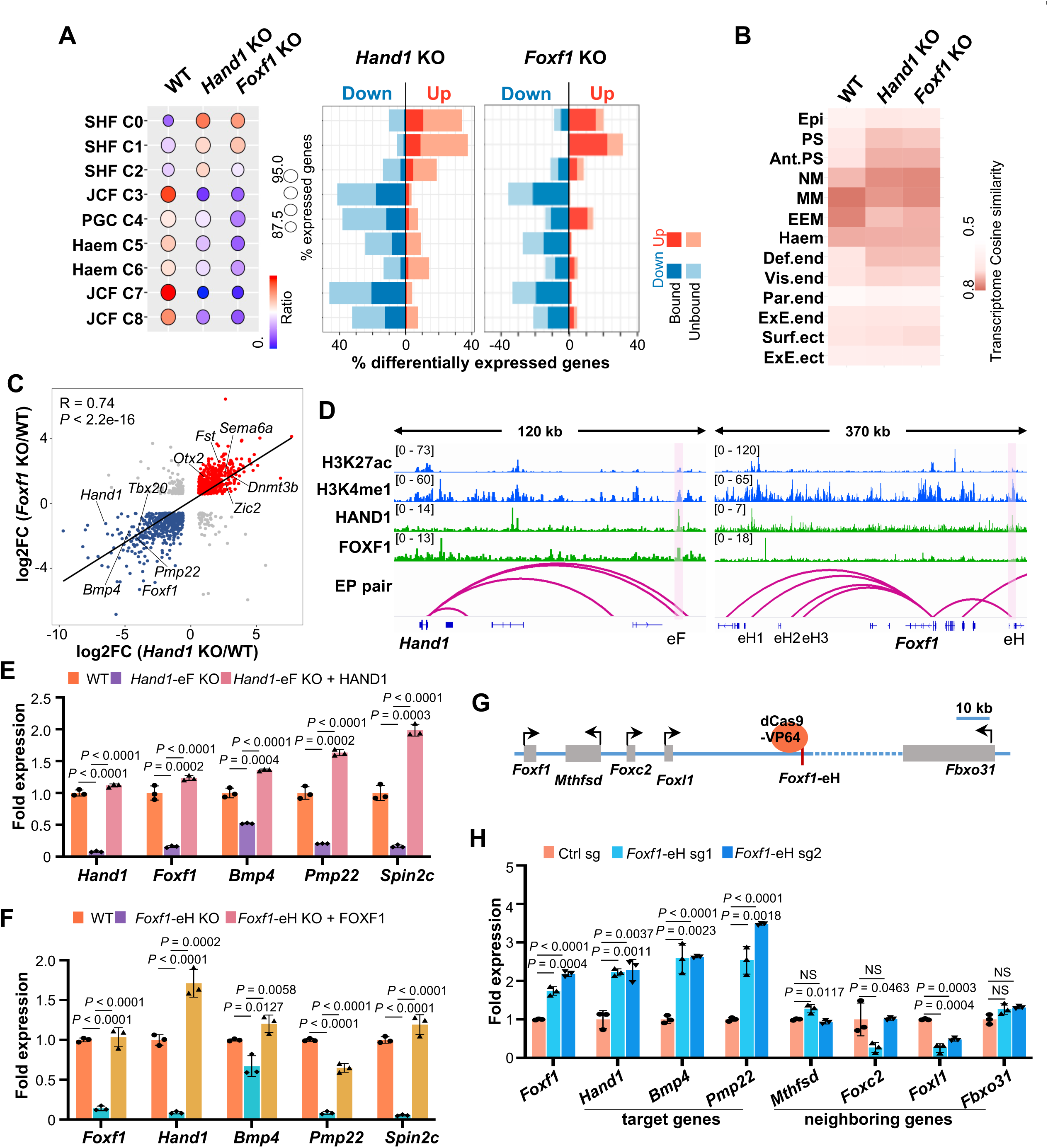
HAND1 and FOXF1 are mutually regulated and required for the expression of the JCF specific genes. **(A)** Comparison of the genes affected by *Hand1* KO or *Foxf1* KO with cluster specific genes of E7.0 mesodermal cells. Dot plot showing that absolute (dot size) and relative (dot color) ratio of the cluster specific genes in WT, *Hand1* KO or *Foxf1* KO MES cells. >90% cluster specific genes were expressed in MES. Higher enrichment in C0-2 and lower enrichment in other clusters can be observed in *Hand1* KO or *Foxf1* KO MES cells. Bar plots showing the up- or down-regulated genes in *Hand1* KO or *Foxf1* KO MES cells. Lengths of bars representing the percentage of cluster specific genes which were up- or down-regulated. Dark colors indicating direct targets of *Hand1* (left) and *Foxf1* (right). **(B)** Heatmap showing the transcriptomic similarity of MES cells and cell types of E7.0 mouse embryo. MES transcriptome data were generated using bulk RNA-seq. Cell-type specific transcriptomes of E7.0 mice were determined as the average of single-cell transcriptomes from each cell type. The gene set for comparison was defined as the collection of top 50 marker genes of each E7.0 cell type. Cosine similarity metric was used. Ant.PS, anterior primitive streak. Haem, haematoendothelial progenitors. Def.end, def.endoderm. Vis.end, visceral endoderm. Par.end, parietal endoderm. ExE.end, ExE endoderm. Surf.ect, surface ectoderm. ExE.ect, ExE ectoderm. **(C)** Scatter plot showing gene expression FCs after *Hand1* KO or *Foxf1* KO. Dots representing genes with FC > 1.5 and adjusted *P* (*P*adj)-value < 1e-5 (Wald tests). Red and blue colors indicating genes co-activated/inhibited in *Hand1* KO and *Foxf1* KO cells. Correlation coefficient of FCs between *Hand1* KO and *Foxf1* KO is 0.74, *P*-value < 2.2e-16, *t*-test. **(D)** Representative genome browser snapshots showing the localization of H3K27ac, H3K4me1, HAND1 and FOXF1 at the *Hand1* and *Foxf1* loci. The FOXF1-binding *Hand1* enhancer (*Hand1*- eF) and the HAND1-binding *Foxf1* enhancer (*Foxf1*-eH), for enhancer KO experiments, are highlighted. **(E-F)** RT-qPCR showing that the reduction in RNA levels of the EEM marker genes after *Hand1*- eF KO can be rescued by HAND1 OE (e), and after *Foxf1*-eH KO can be rescued by FOXF1 OE. (G) Cartoon illustrating target *Foxf1*-eH of by dCas9-VP64 and the genes in the vicinity of *Foxf1*- eH. (H) RT-qPCR showing that the substantial increase in RNA levels of *Foxf1* and the FOXF1 target genes, but not the *Foxf1*-eH neighbouring genes in *Foxf1*-eH CRISPRa cells. Data are the mean ± standard error of the mean from three independent experiments. Two-tailed unpaired Student’s *t*-test was performed.

### Mutual regulation between HAND1 and FOXF1 in driving cardiac specific gene expression

We found that around 50% of the dysregulated genes in *Foxf1* KO MES cells were also dysregulated in *Hand1* KO MES cells, suggesting their synergistic function in transcriptional regulation (Figure 4C). For example, the key JCF specific genes, including *Hand1*, *Foxf1*, *Bmp4*, *Tbx20* and *Pmp22* were significantly down-regulated, while the epiblast genes *Dnmt3b*, *Sema6a* and *Fst*, and the NM specific genes *Otx2* and *Zic2* were substantially up-regulated in both KO cells (Figures 4C and S6E). Depletion of *Hand1* blocked the activation of *Foxf1* during cardiac progenitor differentiation, and vice versa, while *Hand1* overexpression was able to activate *Foxf1* and the other JCF specific genes, and vice verse (Figures S7A-D).

The functional relevance between HAND1 and FOXF1 in target gene regulation could be attribuited to the enrichment of HAND1 at the *Foxf1* enhancers, and vice versa (Figure 4D). We then used CRISPR-Cas9 technology to delete the putative enhancers of *Hand1* and *Foxf1* (Figures S7E-F). Indeed, deletion of the HAND1-bound *Foxf1* enhancer (*Foxf1*-eH) abolished the activation of *Foxf1*; deletion of the FOXF1-bound *Hand1* enhancer (*Hand1*-eF) also significantly reduced the expression of *Hand1* during cardiac differentiation (Figures 4E and 4F). Importantly, the induction of the JCF specific genes *Bmp4*, *Pmp22* and *Spin2c* was severely impaired after deletion of either *Foxf1*-eH or *Hand1*-eF. Overexpression of HAND1 in the *Hand1*-eF KO cells and FOXF1 in the *Foxf1*-eH KO cells were able to rescue the levels of these JCF specific genes during cardiac differentiation. To further investigate the role of *Foxf1*-eH in *Foxf1* expression, we performed CRISPR activation (CRISPRa) assay to activate *Foxf1*-eH (Figures 4G). CRISPRa of *Foxf1*-eH led to a specific increase the expression of *Foxf1* and its downstream target genes, but not its neighboring genes (Figure 4H). Together, our in vitro experimental data indicated that mutual regulation between HAND1 and FOXF1 could play a key role in activation of JCF specific genes.

### *Hand1* KO leads to MES overproliferation but cell death after exiting from the MES status

Meta analysis demonstrated that HAND1 occupied the early Cluster 3 specific DAEs, while FOXF1 tended to show more specific enrichment at the late cardiac specific Clusters 7 and 8 enhancers (Figures 3C and 5A). We also noticed that the *Hand1* KO MES colonies were evidently much larger than those of WT control and *Foxf1* KO, though the same number of EB cells were seeded at equal density for MES differentiation (Figure 5B). While *Foxf1* KO barely affected cell proliferation rate, the count of the *Hand1* KO cells was substantially increased when the cells were differentiated from EB towards MES state (Figure 5C). The *Hand1* KO cells gradually lost viability upon *in vitro* cardiac lineage induction. Consistent with the increased proliferation rate observed, gene ontology (GO) analysis revealed that the genes involved in negative regulation of proliferation and positive regulation of cell migration were specifically down-regulated in *Hand1* KO, but not *Foxf1* KO, MES cells (Figures 5D and S7G).

**Figure 5:**
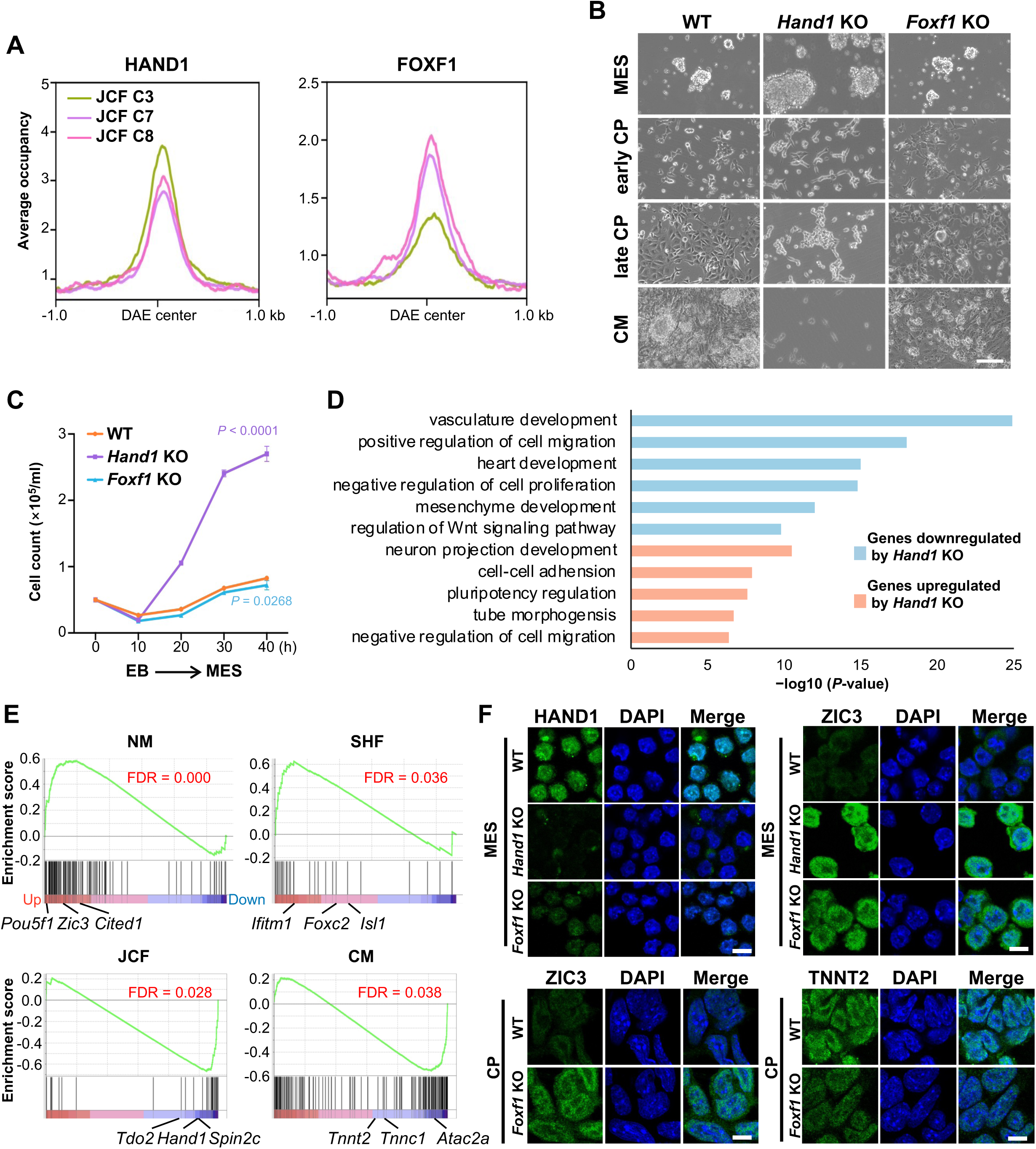
HAND1 and FOXF1 are required for the cardiac lineage specification. (A) Meta plot analysis showing the occupancies of HAND1 (left) and FOXF1 (right) at the C3/7/8 JCF specific DAEs from Figure 2D. (B) Bright field microscopic images of WT, *Hand1* KO and *Foxf1* KO cells at MES, early CP, late CP and CM stages. Scale bar: 100 μm. (C) Growth curve showing numbers of WT, *Hand1* KO and *Foxf1* KO cells counted at different time intervals during EB to MES differentiation. Data are the mean ± standard error of the mean from three independent experiments. Two-tailed unpaired Student’s *t*-test was performed. (D) Enriched GO terms of the genes up- or down-regulated after *Hand1* KO. One-sided Fisher’s Exact test with Benjamini-Hochberg multiple testing correction was performed. (E) GSEA showing the distribution of the marker genes within the ranking of *Foxf1* KO affected genes of *in vitro* differentiated CP cells (FDR calculated by permutation tests). *Foxf1* KO affected genes were ranked from up- (red) to down-regulation (blue). (F) Immunofluorescence staining of HAND1, ZIC3, and TNNT2 in WT and *Hand1/Foxf1* KO lines at MES/CP stage. Scale bar: 10 μm.

The *Foxf1* KO MES colonies derived from MES differentiation appeared phenotypically normal, but were not able to further differentiate into beating CMs (Figure 5B and Supplementary Video 1). The expression of the mature CM markers *Tnnt2*, *Myh6* and *Myh7* were also substantially lower after *Foxf1* KO (Figure S7H). To further examine the function of FOXF1 in cardiac progenitor specification, we performed RNA-seq analysis in control and the CP cells derived *Foxf1* KO mESCs. Gene Set Enrichment Analysis (GSEA) revealed the expression levels of the NM specific genes (like *Zic3*, *Pou5f1* and *Cited1*) and the SHF specific genes (like *Isl1*, *Ifitm1* and *Foxc2*) were remarkably up-regulated, while the expression levels of the JCF specific genes (like *Hand1*, *Spin2c* and *Tdo2*) and the early CM specific genes (like *Acta2*, *Tnnc1* and *Tnnt2*) were significantly reduced (Figure 5E). Thus, our data further supported the specific and synergistic roles of HAND1 and FOXF1 in JCF cardiac progenitor specification. In addition, we performed IF staining of mesodermal (ZIC3), JCF (HAND1) and cardiac markers (TNNT2), followed by cell quantification (Figure 5F). Results indicate that *Hand1* and *Foxf1* knockout leads to reduced commitment to the JCF lineage, evidenced by the loss of *Hand1* expression, accumulation of undifferentiated ZIC3+ mesoderm, and impaired cardiomyocyte formation (TNNT2+), consistent with the up-regulation of JCF lineage specific genes and the down-regulation of SHF lineage specific genes. These results suggest that HAND1 and FOXF1 may cooperatively regulate early cardiac lineage specification by promoting JCF-associated gene expression and suppressing alternative mesodermal programs.

### Genetic loss of *Hand1* blocks the specification of mesoderm along JCF

To further invesigate the roles of HAND1 in JCF specification *in vivo*, we generated floxed allele of *Hand1* by inserting loxP sites flanking the exon 1 of the *Hand1* gene. Genetic crosses of the *Hand1*^fl/fl^ mice with *Mesp1*-*Cre* mice[7] allowed specific deletion of *Hand1* in mesodermal cells (Figure S8A). Consistently, *Foxf1* is also co-expressed in HAND1-positive EEM cells and its expression was also drastically down-regulated in MESP1-CRE driven *Hand1* conditional KO (*Hand1* CKO) embryos (Figure 6A). Mesodermal deletion of *Hand1* generally led to smaller embryos at E7.0 and also later stages (Figure S8B). Data suggest that by E8.5 when heart looping initiate in control group (14/17), the hearts of *Hand1* CKO embryos (3/3) still demonstrate a linear tube morphology (Figure S8C). By E9.5 when atrium and ventricle become distinct in WT embryos, heart looping of *Hand1* CKO embryos is abnormal (Figure 6B). The *Hand1* CKO embryos appeared to die by E9.5 due to embryonic turning failure and heart looping abnormality, mirroring the previously reported phenotype of *Hand1* KO mice [18, 19].

**Figure 6:**
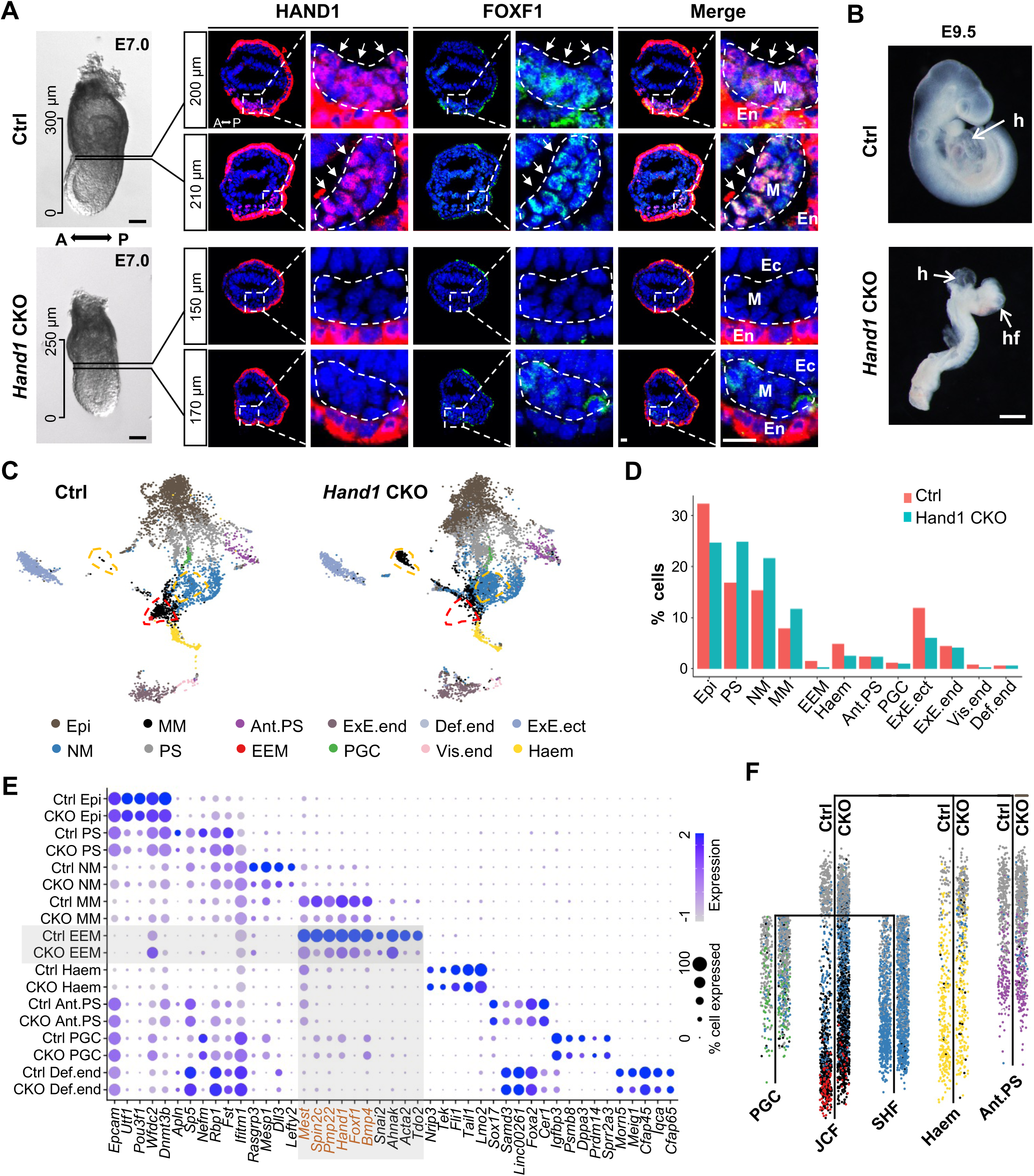
*Hand1* KO in mesodermal cells blocks JCF specification in mouse embryos. **(A)** Immunofluorescence staining showing a substantial reduction in protein levels of HAND1 and FOXF1 at the embryonic-extraembryonic boundary transverse sections (indicated by dashed lines and arrows) of E7.0 *Hand1* CKO embryos. Bright filed images (lateral view) of corresponding embryos are shown on the left side. The distance for each section to the distal tip of the embryo is labelled at the upper right corner for each image. Scale bars: 100 μm (bright field images, left) and 20 μm (transverse sections, right). **(B)** The bright field images of E9.5 Control (Ctrl) and *Hand1* CKO mouse embryos. The arrows indicating the embryonic heart (h) and head folds (hf). Scale bar: 500 μm. **(C)** UMAP layout of integrated E7.0 mouse embryo scRNA-seq data. Integration was performed using the scRNA-seq data of Ctrl and *Hand1* CKO mice. Colors indicating cell types. Dashed lines emphasizing cells with increased (red) or decreased (yellow) numbers in *Hand1* CKO mice. **(D)** Bar plot showing the percentage of each cell type of Ctrl and *Hand1* CKO mice. **(E)** Dot plot showing marker gene expression in Ctrl and *Hand1* CKO mice. Red highlighting markers of MM and EEM cell type. **(F)** URD inferred trajectory tree revealing the developmental hierarchy of E7.0 mesoderm cells (Ctrl and *Hand1* CKO snRNA-seq), colored by cell types. Ctrl and *Hand1* CKO cells are distributed on both sides of the URD tree trunk, with Ctrl cells on the left and *Hand1* CKO cells on the right.

To assess how *Hand1* loss affects early mesoderm development *in vivo*, we analyzed the single-cell transcriptomics of control and *Hand1* CKO embryos at E7.0. Integrated analysis indicated the loss of EEM cells, but the abnormal accumulation of primitive streak (PS), NM and MM cells in *Hand1* CKO embryos (Figures 6C and 6D), which suggested that specific deletion of *Hand1* in mesodermal cells strongly affected early mesoderm differentiation. Detailed analysis regarding the percentage of each cell type revealed the specific reduction of cell numbers from EEM, hematoendothelial progenitors, and ExE ectoderm cell clusters in *Hand1* CKO embryos (Figure 6D). To further support this finding, we performed immunofluorescence staining of the EEM marker *Vim*, which revealed a marked reduction of *Vim*+ cells in *Hand1* CKO embryos compared to controls, indicating impaired progression toward the JCF lineage (Figure S8D).We then compared the gene expression pattern for each cell cluster and calculated the significantly affected genes between Control and *Hand1* CKO embryos (Figure 6E). Consistently, EEM cell cluster was the most affected clusters upon *Hand1* loss (Figures 6E and S8E). Although the percentages of hematoendothelial progenitors and ExE ectoderm cells were reduced, the expression of both cell type specific marker genes did not seem affected drastically. To further illustrate the developmental progression of the mesodermal lineage, we performed paired URD lineage inference analysis[20], further confirming the specific development block of NM specification towards EEM in JCF (Figure 6F). We also performed label transfer analysis to identify JCF and SHF progenitor cells from the E7.0 scRNA-seq data (Figure S9A). Our analysis showed that the fraction of JCF progenitors increased by over 2-fold, whereas the fraction of SHF progenitors remained unchanged (Figures S9B and S9C), supporting that *Hand1* deletion driven by MESP1- CRE led to accumulated JCF progenitor cells and blocked the JCF direction of mesoderm differentiation.

The abnormal accumulation of PS, NM and MM cells in *Hand1* CKO embryos appeared to be consistent with the phenotypes observed in *in vitro* mesoderm differentiation of *Hand1* KO mESCs (Figures 4B and 5B). Notably, the genes involved in negative regulation of proliferation and down- regulated in *Hand1* KO MESs were also reduced in the JCF, but not SHF, trjectory of E7.0 *Hand1* CKO embryos (Figure 7A). In addition, cell migration related genes were also affected in *Hand1*- depleted MES and embryos. To further validate this phenotype, we performed sequential DAPI staining on cryo-sectioned E7.0 control and *Hand1* CKO embryos, followed by cellular segmentation and cell density measurement (Figures 7B and 7C). The analysis revealed that the mesoderm cells near the extraembryonic region in *Hand1* CKO embryos were more compacted (Figures 7B’’, 7C’’, and 7D), while the distal region of the *Hand1* CKO embryos showed no obvious difference from the control embryos (Figures 7B’’’ and 7C’’’). In addition, reduced exocoelomic cavity (EC) size and increased number of mesodermal cells in extraembryonic region were also observed in *Hand1* CKO embryos (Figures 7B’ and 7C’). Hematoxylin and eosin (H&E) staining of E7.5 embryos also supported reduced embryo size and accumulated cells at mesodermal regions upon loss of *Hand1* (Figure 7E). These data together establish HAND1 as a factor in promoting the specification of mesodermal cells toward JCF.

**Figure 7:**
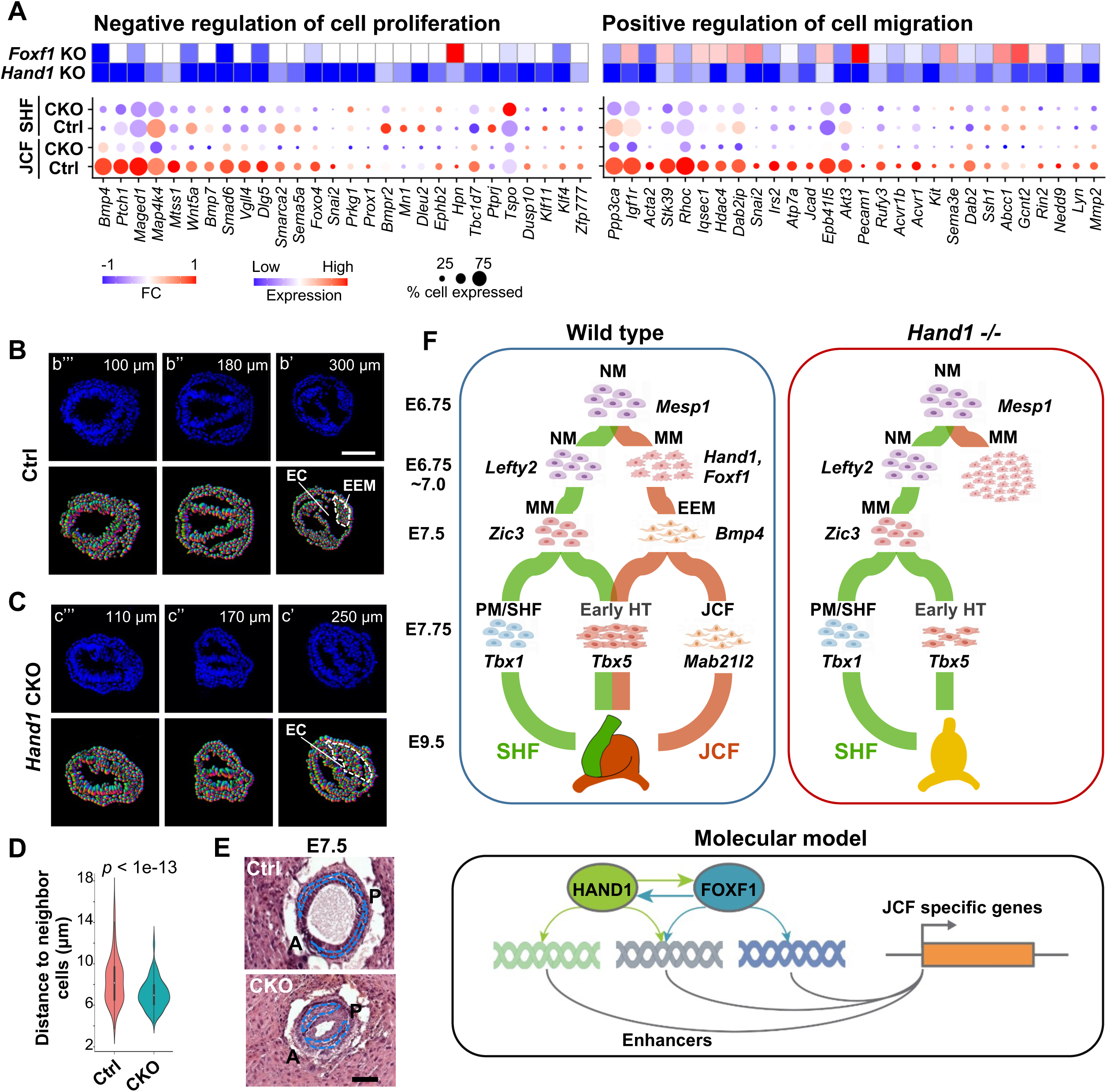
HAND1 is required for exiting from the NM program. **(A)** Co-regulated changes in the genes related to negative regulation of cell proliferation (left) and positive regulation of cell migration (right) *in vivo* and *in vitro*. Heatmap showing the fold changes (FC) of those co-regulated genes after *Hand1* KO and *Foxf1* KO in MES cells *in vitro* (top). Dot plot showing expression levels of those co-regulated genes after *Hand1* KO in the JCF and SHF trajectories *in vivo* (bottom). **(B-C)** DAPI staining in the transverse sections of E7.0 Ctrl (b) and *Hand1* CKO (c) embryos. The distance for each section to the distal tip of the embryo was labelled at the upper right corner for each image. Dashed white lines denoting locations of the exocoelomic cavity (EC) and NM. **(D)** Violin plot showing the distance between neighboring mesoderm cells in the transverse sections of E7.0 Ctrl and *Hand1* CKO embryos. *P*-value was calculated using one-sided Mann-Whitney U test. **(E)** H&E staining showing reduced embryo size and accumulation of mesodermal cells of E7.5 *Hand1* CKO mouse embryos. Scale bars: 100 μm. **(F)** Working model of early cardiac lineage differentiation. Cardiac fate specification initiates around E6.75-7.0 by segregation of the JCF and SHF trajectories. At E7.0, multipotent progenitor cells in JCF lineage present *Hand1*+ mixed mesodermal (MM) status, in contrast to SHF cells showing *Lefty2*+ nascent mesodermal (NM) status. JCF and SHF specifically contribute to JCF and SHF, respectively, while they both contribute to the formation of early heart tube (HT). HAND1 and FOXF1 bind to enhancers, regulate the JCF specific genes, and drive mesodermal differentiation toward JCF direction. *Hand1* deletion led to blocked JCF specification and accumulation of early mesodermal cells.

## Discussion

In this study, we described the transcriptional trajectories and epigenetic landscapes of early cardiac specification event in mouse embryos, and identified a predicted CRN underlying the early mesodermal lineage specification. Mechanistically, we demonstrated that this earliest cardiac JCF specification event was tightly regulated by HAND1 and FOXF1. HAND1 and FOXF1 were mutually regulated, but also performed their respective functions in JCF cell fate determination. *In vitro* differentiation and *in vivo* mouse model analyses indicated that HAND1 was essential for exiting from NM program during cardiac specification. Depletion of FOXF1 impaired the capacity of mesodermal cells in cardiac specification (Figure 7F). In sum, our findings provided new insights into the transcriptional determinants that specify the early cardiac progenitors, paving the way for the identification of potential therapeutic targets for treating congenital heart defects.

Recent studies using scRNA-seq have reported the roadmaps of mammalian embryonic lineage development[12, 21]. Together with studies which focus on cardiac progenitors[4, 6, 22], several models have been proposed to explain the multi-lineage process of heart formation. In this study, we observed a clear divergence between the JCF and SHF trajectories around E7.5-E7.75, and performed forward-backward tracing to reconstruct their developmental paths. Although cells were assigned to distinct trajectories based on dominant fate probabilities, fate divergence analysis revealed that JCF and SHF likely originate from a common progenitor pool prior to E7.0. The two trajectories are consistent with the previous clonal analysis and also the HAND1+ cells lineage tracing by Zhang et al. [23]. Importantly, we here identified specific trajectory-related gene cohorts throughout the whole process, shedding light on the molecular mechanisms by which HAND1 is crucial for promoting the exit of NM program during the early JCF specification process.

MESP1 serves as a master regulator in the establishment of the cardiac lineage. An analysis of previously published MESP1-regulated genes revealed that GATA4, HAND1 and FOXF1 were directly controlled by MESP1. GATA4 was rapidly induced by MESP1 within 12 hours, while HAND1 and FOXF1 were activated after 24 hours of MESP1 induction. On a pseudotime scale, we noticed a sequential-temporal expression pattern of GATA4, HAND1 and FOXF1 in JCF. Our transcriptional and epigenomic analyses suggested a sequential activation model of MESP1- GATA4-HAND1-FOXF1. However, it should be noted that MESP1 and GATA4 were activated in both JCF and SHF. Thus, additional regulation must exist and instructs the process of JCF-SHF lineage segregation. It has been reported that BMP signaling activated the expression of *Hand1* during heart formation [24]. Since Bmp4 displayed higher specificity for JCF, it might explain the activation of *Hand1*, and subsequent *Foxf1* activation in JCF, but not the SHF. Thus, HAND1 and FOXF1 might be subject to feed-forward activation by MESP1 and GATA4 in concert with BMP signalling, thereby promoting JCF cardiac specification. Future studies are worthwhile to further investigate how TFs work together with signaling pathways to promote cardiac lineage segregation.

We further demonstrated the synergistic roles of HAND1 and FOXF1 in early cardiac specification. *Foxf1*-/- mice were not able to survive beyond E10.0, with incomplete separation of splanchnic and somatic mesoderm, leading to abnormal coelom formation. Interestingly, FOXF1 marked the mesothelium lining (a monolayer epithelial lining of the exocoelomic cavity), the anterior distal part of which extends to the presumptive cardiac mesoderm [25]. Such an expression pattern is coincident with that of HAND1 in JCF [4]. The role of FoxF is also essential for the early cardiac development in the ascidian Ciona intestinalis [26]. During the early heart development, HAND1 demonstrates multi-facet functions and contribute to the formation of FHF [6], heart loop [19], and outflow tract [27]. A recent study suggests that *Hand1* knockout biases progenitor cells toward SHF lineage, affecting the expression of cardiomyocyte markers and delaying differentiation onset in human pluripotent stem cells [28]. Our studies showed that FOXF1 was indispensable for the specification of mesoderm toward early cardiac progenitors, through binding the JCF progenitor specific enhancers for their later activation. Together, the combined analyses on the developmental route of CMs and their transcriptional determinants will further enhance our understanding of the etiology behind congenital heart defects, ultimately providing insights into potential regenerative strategies for heart disease treatment.

## Acknowledgements

The authors are grateful to the Lin and Luo lab members for helpful discussion of this study. The authors thank Profs. Shaorong Gao, Xinhua Lin, and Yaofeng Zhao for critical discussion and suggestions to this study. The authors thank Min Wang for assistance in generating the Hand1 KO cell lines. Studies in this manuscript were supported by funds provided by National Key R&D Program of China (2018YFA0800100 and 2018YFA0800101 to C.L.; 2018YFA0800103 to Z.L. and P.X.), the National Natural Science Foundation of China (32030017 to C.L.; 32100529 to P.X.).

## Author contributions

C.L., Z.L., P.X., and N.J. designed the research. X.J., Q.P., X.Y., and Y.Z. performed most the experiments and analyzed the data. P.X., J.H., C.W., W.Z., S.S., and Y.Y. analyzed transcriptomic and epigenomic data. Z.C., K.F., and H.F. assisted with cell line generation. W.F., K.G., and S.Z. performed the breeding of mouse lines. T.Z., Z.Y., K.W., W.X, Z.L., and C.L. provided the resources. Z.L., C.L., and P.X. wrote the manuscript. C.L. and Z.L. supervised the project.

## Declaration of interests

The authors declare that they have no conflicts of interest.

## STAR Methods

### Mice

All mouse experiments were approved by the Animal Care and Use Committee at Southeast University, and performed in accordance with institutional guidelines. Mice were housed in cages under SPF conditions and had free access to water and food. *Hand1*^fl/fl^ mice were produced by Cyagen with loxP sites inserted into the regions surrounding exon 1 in the *Hand1* locus. *Hand1*^fl/+^; *Mesp1*-*Cre* mice were generated by crossing *Hand1*^fl/fl^ mice with *Mesp1*-*Cre* mice [32].

The *Hand1*^fl/fl^ or *Hand1*^fl/+^ female mice caged with male *Hand1*^fl/+^; *Mesp1*-*Cre* mice. Females were screened for vaginal plugs following morning (E0.5). To obtain post-implantation embryos, female mice at 7.0 or 9.5 days post-coitum (d.p.c.) were executed and the uteri were dissected and transferred to a petri dish with PBS. For 7.0 d.p.c embryos, each decidua was carefully freed from the uterine muscle layers using properly sharpened forceps. Then the embryos were carefully separated from decidua. Reichart’s membrane and the ectoplacental cone were also removed from the 7.0 d.p.c embryos. The uteri surrounding the 9.5 d.p.c embryos were cut open with a small incision, the embryos were genetly squeezed out, and then the amniotic membranes were removed. Images of the embryo were acquired on a stereo microscope (Mshot, MZ62; Olympus, SC180).

Genomic DNA for mouse genotyping was obtained from mouse tail biopsies. Genotyping of mouse embryos was performed with genomic DNA, which was obtained by collecting extra- embryonic regions subsequently digestion using Mouse Direct PCR Kit (Bimake, B40013). PCR reactions were used to detect the *Cre* transgene and the *Hand1* loxP site. Thermal cycle reactions were as follows: 3 min at 94 °C, 35 cycles of 30 s at 94 °C, 35 s at 60 °C, 45 s at 72 °C and a final 5 min extension at 72 °C. Primer sequences used in this study are listed in Supplementary Table 3.

### Antibodies

Antibody against HAND1 (sc-390376) (WB: 1:1000, IF for embryos: 1:100, IF for cells: 1:200) was purchased from Santa Cruz. Antibody against FOXF1 (Abclonal, A13017) (WB: 1:2000, IF for embryos: 1:100, IF for cells: 1:500) was purchased from Abclonal. Antibody against HA (ChIP: 3 μg) was generated in house. Antibody against TUBULIN (66031-1-Ig) (WB: 1:50, 000) was purchased from Proteintech. Goat anti-mouse IgG Alexa Fluor 488(A-11001) and goat anti-rabbit IgG Alexa Fluor 546 (A-11035) were purchased from Thermo Fisher Scientific.

### Mouse ESC culture

Mouse E14 ESCs were maintained in DMEM (Hyclone, SH30243.01) supplemented with 15% FBS, 1 × nonessential amino acids (STEMCELL, 07600), 1 × GlutaMAX (Gibco, 35050-061), 1 × penicillin streptomycin solution (Sangon Biotech, E607011-0500), b-mercaptoethanol (ALDRICH), 0.1 μg/ml leukemia inhibitory factor (Novoprotein, C690), 3 μM CHIR99021 and 1 μM PD0325901 in gelatin-coated plates at 37 °C under 5% CO2.

### CRISPR-Cas9 guided KO

SgRNAs targeting *Hand1*, *Foxf1* and their enhancer sites were cloned into lentiCRISPR v2. These constructs were transfected into 70% confluent 293T cells together with 6 μg of psPAX2 packaging plasmids and 2 μg of pMD2.G envelope plasmids using Highgene (Abclonal, RM09014). The media was half-replaced with fresh DMEM supplemented with 10% FBS 6 h after transfection. The lentiviral supernatants were harvested at 24 h, 48 h, and 72 h post-transfection, filtered through 0.45 μm filters, and concentrated at 50,000 × g for 0.5 h. Mouse E14 ESCs were infected with concentrated lentiviral particles with polybrene (Sigma) at the concentration of 8 μg/ml. 2 μg/ml puromycin was used for selection for 48 h and individual colonies were picked and expanded in 48-well plates. The clones were screened with genomic PCR, and confirmed by TA cloning and Sanger sequencing. Oligo sequences used in this study are listed in Supplementary Table 3.

### *In vitro* cardiac differentiation

Mouse E14 ESCs were differentiated into embryoid bodies (EBs) at a density of 75,000 cells/ml in 6 cm dishes for 48 h in serum-free media (DMEM (Hyclone, SH30243.01), DMEM/F12 (1:1) (Gibco, 11320-033), 0.05% BSA (Sigma, A1933), 1 × GlutaMax (Gibco, 35050-061), B27 supplement (Gibco, 12587010), N2 supplement (Gibco, 17502048), supplemented with 50 mg/ml ascorbic acid (Alfa Aesar, 50-81-7) and 4.5×10^-4^ M monothioglycerol). EBs were dissociated into single cells and re-aggregated as MES cells for 40 h at 50,000 cells/ml in the presence of 5 ng/mL human VEGF (Novoprotein, C083) and 10 ng/ml human Activin A (Peprotech, 96-120-14-10) and 0.3 ng/ml human BMP4 (R&D, 5020-BP-010). MES cells were dissociated and plated as a monolayer in gelatin-coated 12-well plate at 50,000 cells/ml in StemPro-34 (Gibco, 10639011) supplemented with 5 ng/mL VEGF (Novoprotein, C083), 10 ng/mL human basic FGF (Gibco, PMG0035) and 25 ng/mL FGF10 (R&D, 345-FG-250). CP cells were harvested after differentiation for 32 h. Contracting CMs can be observed after another 5 days. Differentiation stages were confirmed by the expression of marker genes.

### Immunofluorescence

Mouse Embryos: Embryos were fixed in 4% PFA, then rinsed in PBS and incubated in 30% sucrose at 4 °C until sinking. Embryos were then embedded in OCT medium and snap-frozen in liquid nitrogen and stored at −80 °C. Sections were cut at a thickness of 10 μm. To perform IF, embryo sections were washed three times for 5 min each, then antigen repair was performed using citrate. The samples were permeated for 40 min with 0.3% TritonX-100 followed by washing three times with PBS for 5 min each. The samples were then blocked with ReadyProbes 2.5% Normal Goat Serum (Thermo) for 1 h at room temperature. The samples were incubated overnight at 4 °C with primary antibodies followed by washing three times with PBS for 10 min each. The samples were incubated with secondary antibodies for 1 h at room temperature, then washed three times with PBS for 10 min each. The samples were then incubated with DAPI at room temperature for 10 min followed by washing one time with PBS for 5 min. After the final wash, the samples were mounted with mounting buffer. Images were captured using a Zeiss 700 laser confocal microscope.

MES and CP cells were dissociated into single cells and fixed on 12-well plate cell slide with 4% PFA at room temperature for 15 min. The fixed cells were washed 3 times with PBS and permeabilized with 0.2% Triton X-100 for 5 min at room temperature. The cells were then blocked with PBS containing 2% BSA and 0.3% TritonX-100 for 1 h at room temperature. Appropriate dilution of primary antibody was added and cells were incubated overnight at 4 °C. After washing with PBS for 3 times, the cells were stained with secondary antibodies and DAPI for 1 h at room temperature followed by mounting on slides. Images were captured using a Zeiss 700 laser confocal microscope.

### H&E staining

Whole E7.5 embryos were fixed overnight with 4% PFA, embedded vertically in clean paraffin, then sliced to obtain 7 µm paraffin sections. Standard hematoxylin and eosin (H&E) staining methods were used to stain the sections. The images of H&E staining were obtained on a microscope (Olympus, IX73).

### Western blotting

Cells were washed twice in ice-cold PBS and incubated in 1 × SDS lysis buffer for 15 min at 95 ℃. Lysates were pre-cleared by maximum speed centrifugation for 3 min, then separated by SDS-PAGE and transferred to polyvinylidene fluoride (PVDF) membrane. 5% non-fat dry milk was used for blocking and primary antibody was added. The membrane was incubated overnight at 4 °C, then washed, incubated with secondary antibody for 1 h at room temperature. ECL substrate was used for imaging by autoradiography.

### Quantitative RT-PCR and bulk RNA-seq

Total RNA was extracted from cells using RNA isolater Total RNA Extraction Reagent (Vazyme). 500 ng of total RNA was used to synthesize cDNA using ABScript II RT Mix (Abclonal, RK20403). Resultant cDNA was diluted in water and 12.5 ng cDNA was used in each qRT-PCR reaction. Reactions were run on CFX96 (Bio-Rad) using 2 × Universal SYBR Green Fast qPCR Mix (Abclonal, RK21203). The relative expression levels of genes of interest were normalized to the housekeeping gene *Actin*. Each experiment contains at least three biological replicates. Primer sequences used in this study are listed in Supplementary Table 3. For RNA-seq, 1 × 10^6^ MES or CP cells were harvested and RNA extraction was performed using Rneasy mini plus kit (Qiagen). 1 μg of total RNA was used for the construction of sequencing libraries and sequencing.

### snRNA-seq

To prepare single cells for snRNA-seq, embryos at E7.0 were dissected in cold sterile 1 × PBS without Ca^2+^, Mg^2+^ under a stereo microscope. Embryos were staged based on their morphology. The Reichert’s membrane and ectoplacental cone were removed, and genotypic identification of embryos was carried out. Embryos were placed into a 1.5 ml microfuge tube and digested into single cells using TrypLETM Express (Thermo) at 37 °C. Single-cell RNA sequencing was performed on single-cell suspensions using 10 × Genomics.

### 10 × Multiome library preparation and high throughput sequencing

Six embryos staged at E7.0 were collected and washed with cooled 0.5% BSA/PBS for twice. Sufficient embryos were pooled and then subjected to 200 μL TrypLE for cell dissociation for 10 min at 37 °C with frequent gentle mixture. Single-cell suspension of embryos were then quenched and washed with 0.5% BSA/PBS, and finally filtered using 40 μm Flowmi cell strainer. The acquired single cell suspension were then subjected to nuclei isolation, library preparation by following the manufacturer’s instruction. The library was sequenced on Novaseq 6000 platform with recommended sequencing depths and read lengths.

### ChIP, ChIP-seq library preparation

5 × e^6^ cells were crosslinked with 1% formaldehyde for 10 min and quenched with 0.125 M glycine for 5 min at room temperature. Fixed cells were incubated in 0.5 ml lysis buffer (50 mM Tris-HCl [pH 8.0], 1 % SDS, 5 mM EDTA) for 10 mins on ice. After adding 1 ml dilution buffer (20 mM Tris-HCl [pH 8.0], 150 mM NaCl, 2 mM EDTA, 1% Triton X-100), chromatins were sonicated into 200-800 bp fragments using Bioruptor (Diagenode) and immunoprecipitated with protein A agarose beads and specific antibody at 4 ℃ for 12 h. Immunoprecipitates were washed with Wash Buffer I (20 mM Tris-HCl [pH 8.0], 150 mM NaCl, 2 mM EDTA, 1% Triton X-100, 0.1% SDS), Wash Buffer II (20 mM Tris-HCl [pH 8.0], 500 mM NaCl, 2 mM EDTA, 1% Triton X-100, 0.1% SDS), Wash Buffer III (10 mM Tris-HCl [pH 8.0], 0.25 M LiCl, 1 mM EDTA, 1% deoxycholate, 1% NP-40) and TE, respectively. After the final wash, DNA was eluted and reverse-crosslinked at 65 °C for at least 6 h. DNA was then purified and used for PCR amplification or ChIP-seq library preparation. ChIP-seq libraries were prepared with VAHTS Universal DNA Library Prep Kit for sequencing.

### Data processing and quality control

#### Single cell multi-omics data: sequence alignment

We used Cell Ranger ARC software suite for the analysis of single cell multi-omics (ATAC & Gene Expression) data (https://support.10xgenomics.com/single-cell-multiome-atac-gex/software/pipelines/latest/what-is-cell-ranger-arc). We applied the “cellranger-arc count” function for barcode counting, adapter/primer removal and sequence alignment. The processed reads were aligned to mm10 using the BWA-MEM algorithm. For ATAC sequencing data, BAM files were generated for downstream analysis. For Gene Expression (GEX) data, “cellranger-arc count” generated gene-barcode matrices were used for further analysis. Version of Cell Ranger ARC genomic sequence and gene annotation: refdata-cellranger-arc-mm10-2020-A-2.0.0.

#### snRNA-seq data

snRNA-seq datasets of the current study include: (1) E6.5-8.5 whole mouse embryos; (2) micro- dissected anterior cardiac regions of mouse embryos at early crescent to linear heart tube (∼E7.75- 8.25) stages. (3) GEX part of the single cell multi-omics data of E7.0 mouse embryos. (4) Control and *Hand1* CKO E7.0 mouse embryos.

We used R package ‘Seurat’ for QC and normalization purposes [33]. Cells with abnormal sequencing depth (nFeature_RNA < 2000 or nCount_RNA > 1e5) or with high mitochondrial ratio (percent.mt > 5) were excluded. We used the ‘SCTransform’ to perform normalization, variance stabilization, and regression of cell cycle scores (using the ‘CellCycleScoring’ function). For dataset (1), we selected E7.0 samples, removed cells labelled as ‘doublet’ or ‘stripped’ and regressed out sequencing batches. Doublet removal was performed using R package ‘DoubletFinder’ [34].

#### snATAC-seq data

Bam files of snATAC-seq from multiome data were processed using Snaptools [30] function ‘snap- pre’ to remove low-quality fragments (MAPQ < 30), over-sized fragments (length > 1,000 bp), secondary alignments and PCR duplicates. This function also generates cell-by-bin matrices of variable resolution (bin sizes: 1kb, 5kb, 10kb) for downstream analysis. The range of fragment coverage for bin selection was set between 500 and 20,000.

#### ChIP-seq data analysis

Adapter sequences were removed from Fastq files using trim_galore (v0.6.7) with default parameters. After trimming, ChIP-seq reads were aligned to the mm10 mouse genome using Bowtie2 (v2.3.5.1) [35] with ‘--no-mixed’ and ‘--no-discordant’ parameters. The aligned files were sorted and converted to BAM format using SAMtools (v1.9). BigWig files were subsequently generated using deepTools bamCoverage (v3.5.0) [36], employing CPM normalization and ignoring duplicates. Peak calling was performed with MACS3 (3.0.0a6) [37] for HAND1 (*Q*-value < 1e-5) and FOXF1 (*Q*-value < 1e-10) ChIP-seq data.

#### Tracing the CM, JCF and SHF trajectories

The trajectories of each in Figure 1 were inferred using the Waddington-OT Python package (v1.0.8) [11] with a predefined starting cell set (E8.5 CM cells for tracing the CM trajectory, E7.5 EEM cells in the CM trajectory for tracing JCF trajectory, and E7.75 pharyngeal mesoderm cells in the CM trajectory for tracing the SHF trajectory), and cells of each trajectory are selected if WOT score > 0.0001. To specifically distinguish between JCF and SHF, the difference in the WOT score of the two trajectories was calculated, and if the difference is greater than 0, it belongs to the JCF trajectory and vice versa to the SHF trajectory. WOT is designed for time-series scRNAseq data where the time/stage each single cell is given. At any adjacent time points ti and ti+1, WOT estimates the transition probability of all cells at ti to all cells at ti +1. One can select a cell set of interest at any time point ti infer their ancestors at ti -1 or their descendants at ti +1 by sums of the transition probabilities. The WOT package was used with default parameters as in the Waddington-OT online tutorial (https://broadinstitute.github.io/wot/tutorial/).

#### Signaling pathway enrichment analysis

Potential signaling-activated/inhibited genes of Bmp, Yap, Wnt, Nodal, Notch and Fgf were collected from Peng et al. 2019 [38]. Enrichment of signaling target gene sets was determined by the average expression levels of activated/inhibited genes. To assess dynamic changes along pseudotime, we calculated the smoothed expression trends of these gene sets using LOESS regression. Notably, an increase in the expression of inhibited genes over pseudotime may reflect a gradual reduction in signaling activity or potential compensation by alternative pathways. These interpretations are consistent with the dynamic and context-dependent nature of signaling regulation during early development.

#### Spatial mapping of single cells

Locations of mouse E7.0/7.5 mesodermal cells were inferred by comparison with the GEO-seq data, where each sample represents 5–40 cells with defined spatial locations. Transfer component analysis (TCA) [39] was performed to achieve shared representation of scRNA-seq and GEO-seq samples. Single cells were mapped to GEO-seq locations with the highest correlation coefficients.

#### Identification of pseudotime-dependent genes

Single-cell pseudotime trajectory analysis was performed using R package Monocle 2 (v2.22.0) [40] according to the online tutorials (http://cole-trapnell-lab.github.io/monocle-release/). Monocle object was directly constructed using Monocle implemented new Cell Data Set function from Seurat (v4.1.1) object, and Monocle implemented differentialGeneTest function was used to find highly variable genes for ordering. Based on these, we selected key and specific genes in JCF and SHF trajectory, respectively, and further visualized them in the heatmap using cell bin-by-gene matrix.

#### snRNA-seq data integration and label transfer

This research included data integration for the following datasets (dataset numbers in ‘Data processing and quality control: snRNA-seq data’): (1) and (2) for comparison of the predicted trajectories and JCF cells; (1) and (3) for cell-type annotation of multiome single cells; (1) and (4) for cell-type annotation of Ctrl/*Hand1* CKO single cells. For multiome dataset, integration was performed by two steps: 1) whole embryo integration; 2) according to the predicted cell types of (1), mesoderm specific integration was done by selecting relevant cell types (NM, MM, EEM, Haem, PGC).

The integration and label transfer process follows the pipeline provided by Seurat [33]: https://satijalab.org/seurat/articles/integration_introduction.html. 4,000 genes were selected for integration based on variable gene sets of individual datasets. We selected “SCT” as the normalization method for identification of integration or transfer anchors.

#### Clustering analysis of snRNA-seq data

We performed clustering analysis for cells of the integrated mesodermal lineage. Seurat [33] function “FindNeighbors” and “FindClusters” were applied to the Top 20 principle compoments with the following parameter setting: annoy.metric = “cosine”, resolution = 0.95. This analysis resulted in nine clusters (C0-8), which is also used for snATAC-seq analysis.

#### FHF and SHF gene signature scores

For each gene set, we defined the average z-score normalized expression levels as the signature score per cell. The FHF and SHF gene sets were collected from Soysa et al. 2019 [22] Supplementary Table 1 (FHF: “FHF” genes of tab “Figure1de_n = 21,366 cells”; SHF: “MP” genes of tab “Fig 2ab_n = 2,103 cells”).

#### snATAC-seq data analysis

snATAC-seq bam files were combined for each cluster, C0-8, for identification of accessible elements (AE). AEs were then collected as a basic set of peak annotation. We used Snaptools [30] function ‘add_pmat’ to generate a cell-by-peak matrix (pmat), where the value of each element represent the accessibility per cell per peak. Diffusion map followed by UMAP analysis was performed to generate the snATAC-seq version of 2D data visualization.

Cluster specific AEs were identified using SnapATAC function ‘findDAR’. For analysis of C2-8 specific AEs, we used C1, which represents the least differentiated cell group, as the background cluster. For C0-1, we used cells from all other clusters as background. Cluster specific AEs were provided to HOMER [31] function ‘findMotifsGenome.pl’ for motif analysis, and to deepTools [36] for plotting heatmaps of ChIP-seq data.

#### Definition of enhancer regulated target genes

Enhancer-promoter (EP) pairs were predicted by SnapATAC function ‘predictGenePeakPair’ [30]. This method performs logistic regression using peak accessibility (snATAC-seq) and expression (snRNA-seq) of neighboring genes to identify EP links. Candicate EP pairs need to be less than 50kb apart. Cluster specific target genes were defined as EP-associated genes of cluster specific AEs. Functional enrichment analysis was performed using R package clusterProfiler [41] (‘compareCluster’ function, ontology was selected as ‘BP’).

#### Analysis of TF activity

We analyzed TF activity, defined as the relative accessibility of AEs containing the motif sequence[42] of a TF, using R package chromVAR (https://greenleaflab.github.io/chromVAR/articles/Introduction.html). SnapATAC-generated cell-by-peak matrices were converted to ‘SummarizedExperiment’ format as the input of chromVAR. The TF activity is calculated by the ‘computeDeviations’ function by comparing the accessibility of motif-containing AEs with background peaks with similar GC content. Genomic coordinates of motifs were acquired by the ‘getMatrixSet’ function and R package ‘JASPAR2020’. Synergy scores between TF pairs were computed by the ‘getAnnotationSynergy’ function.

#### Classification of activation modes of gene-enhancer pairs

We performed pseudotime trajectory analysis for E7.0 JCF trajectory (C0, C3, C7 and C8) in multiome using the abovementioned method. To further explore the underlying mesoderm developmental mechanism of enhancer clusters in regulating gene expression, we first built gene-enhancer pairs for preselected clusters based on snATAC-seq and snRNA-seq datasets. Secondly, we partitioned cells from E7.0 mesoderm lineage into pseudotime bins and calculated the cell bin- by- gene matrix of snRNA-seq and the cell bin-by-peak matrix of snATAC-seq, respectively. Finally, we divided activation modes of gene-enhancer pairs into three groups, fast, sync., and slow, according to the order of bins corresponding to their activation time.

#### Inference for URD lineage tree

To reconstruct branching developmental trajectory trees for E7.0 mesoendodermal cells (Ctrl and *Hand1* CKO), we used URD (v1.1.1) [20]. We used all cells assigned as epiblast as the root of the tree, and cells of clusters that contained the most differentiated cell-types at the latest pseudotime as the tips, where the CM branch was divided into JCF and SHF trajectories. By the movement of coordinates, the cells of Ctrl and *Hand1* KO are located on either side of the branch, respectively.

#### CRN construction and in silico KO

The CRNs based on multi-omics (scRNA + snATAC) data was constructed by SCENIC+ (v1.0.1) (ref) (details available via https://scenicplus.readthedocs.io/en/latest/pbmc_multiome_tutorial.html). CRN-based in-silico KO was conducted by SCENIC + module "scenicplus.simulation". In this step the expression level of a TF was set to zero and SCENIC + propagates the effect through CRN in an iterative manner to obtain the perturbed expression matrix. Cell-state transition vectors were generated and projected to the tSNE map using "plot_perturbation_effect_in_embedding" function.

#### Cell nucleus segmentation

We employed the cellpose (V2.1.1) [43]algorithm for segmentation of DAPI staining for embyo transverse sections. The pre-trained CP model (--pretrained_model CP) was used to obtain the masks of images. We used option “--diameter 8” to resize the image to conform to the input parameters of the pre-trained model. We utilized the MorphoLibJ plugin of Fiji to obtain morphological metrics of the segmented nuclei. Finally, we utilized the VAA3D (2.938) [44] software to perform partitioning of the embryo, to identify cells belonging to the endoderm, mesoderm or epiblast.

## Statistics and reproducibility

Numbers of biological replicates, statistical tests and *P*-values are reported in the figure legends. If not mentioned otherwise in the figure legend, statistical significance was determined using two-tailed Student’s *t*-test, provided by GraphPad Prism9 statistical software.

## Data availability

Sequencing (sc/snRNA-seq, snATAC-seq, ChIP-seq, and RNA-seq) data that support the findings of this study have been deposited in the Gene Expression Omnibus under accession numbers GSE245713. Previously published ChIP-seq data that were re-analyzed here are available under accession codes GSE165107 (MESP1, ZIC2 and ZIC3, 2.5-day EB), GSE47085 (HAND1, FLK+ MES cells), GSE52123 (GATA4, E12.5 mouse heart) and GSE69099 (FLI1, ES derived hemogenic endothelium). Previously published scRNA–seq data that were re-analyzed here are available under accession codes E-MTAB-6967 (ArrayExpress, E6.5-8.5 mouse embryos) and E-MTAB-7403 (ArrayExpress, micro-dissected heart-related samples of E7.75-8.5 mouse embryos). Previously published GEO-seq data that were re-analyzed here are available under accession codes GSE171588. All other data supporting the findings of this study are available from the corresponding author on reasonable request.

---

**Figure S1:**
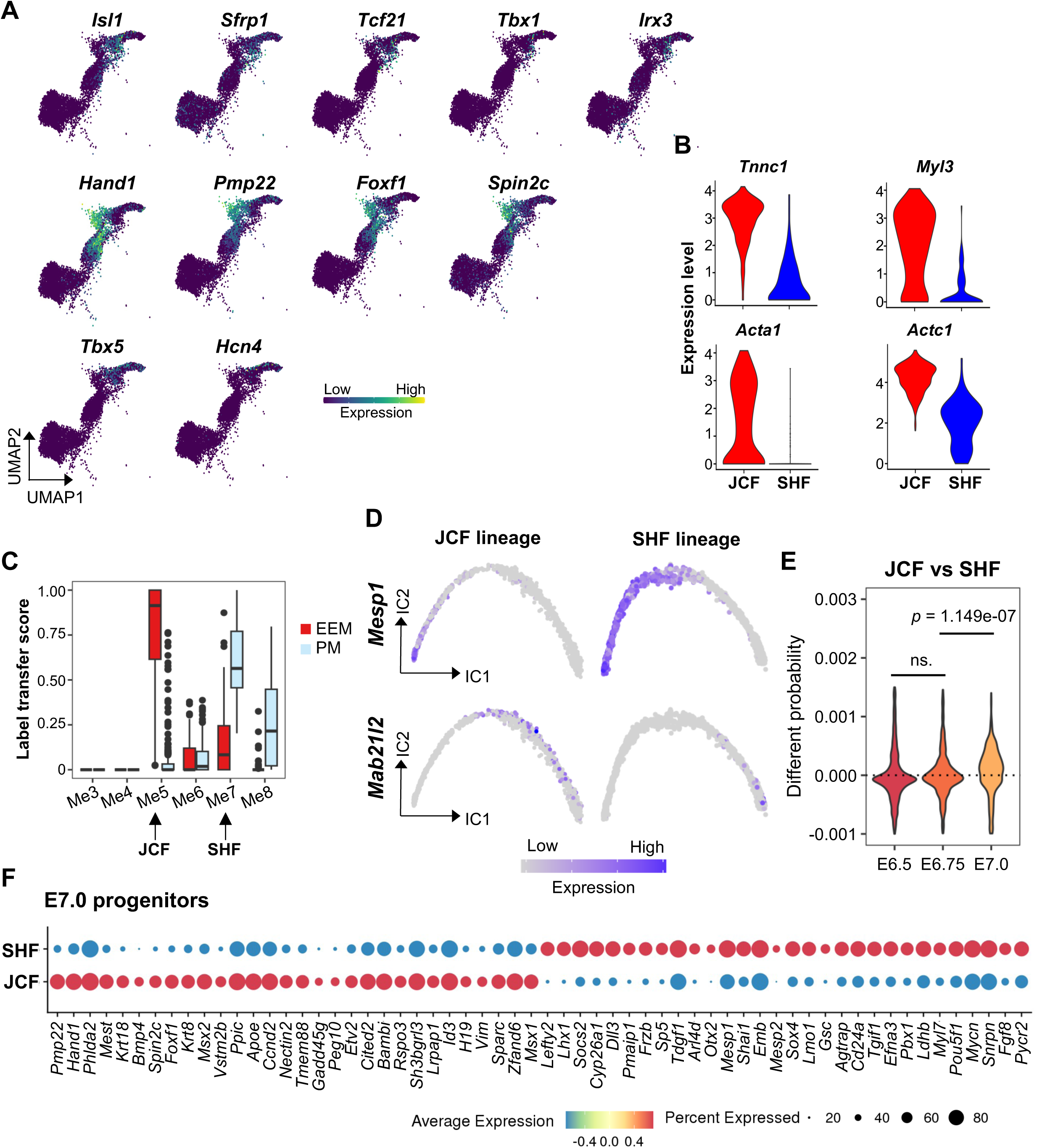
Data integration showing the developmental trajectories of cardiac progenitors **(A)** Gene expression levels of the SHF (upper), JCF (middle) and FHF markers (lower) overlaid on the UMAP layout of the CM trajectory. Only cells belonging to the CM trajectory are shown. **(B)** Violin plots showing the distribution of mature myocardium markers expression levels in E8.5 CM cells of the JCF and SHF trajectories. **(C)** Fraction of labels transferred from Tyser et al. [4] for E7.5 EEM cells and E7.75 PM cells. **(D)** Independent component (IC) layout showing pseudotemporal trajectories for JCF trajectory (left) and SHF trajectory (right) cells, colored by *Mesp1* and *Mab21l2* expression. **(E)** Inference for fate divergence between two trajectories during E6.5-7.0. The violin plots show the differential WOT scores of JCF and SHF lineages. Two-tailed unpaired Student’s *t*-test was performed. **(F)** Marker gene analyses based on JCF/SHF lineages for E7.0 progenitors.

**Figure S2:**
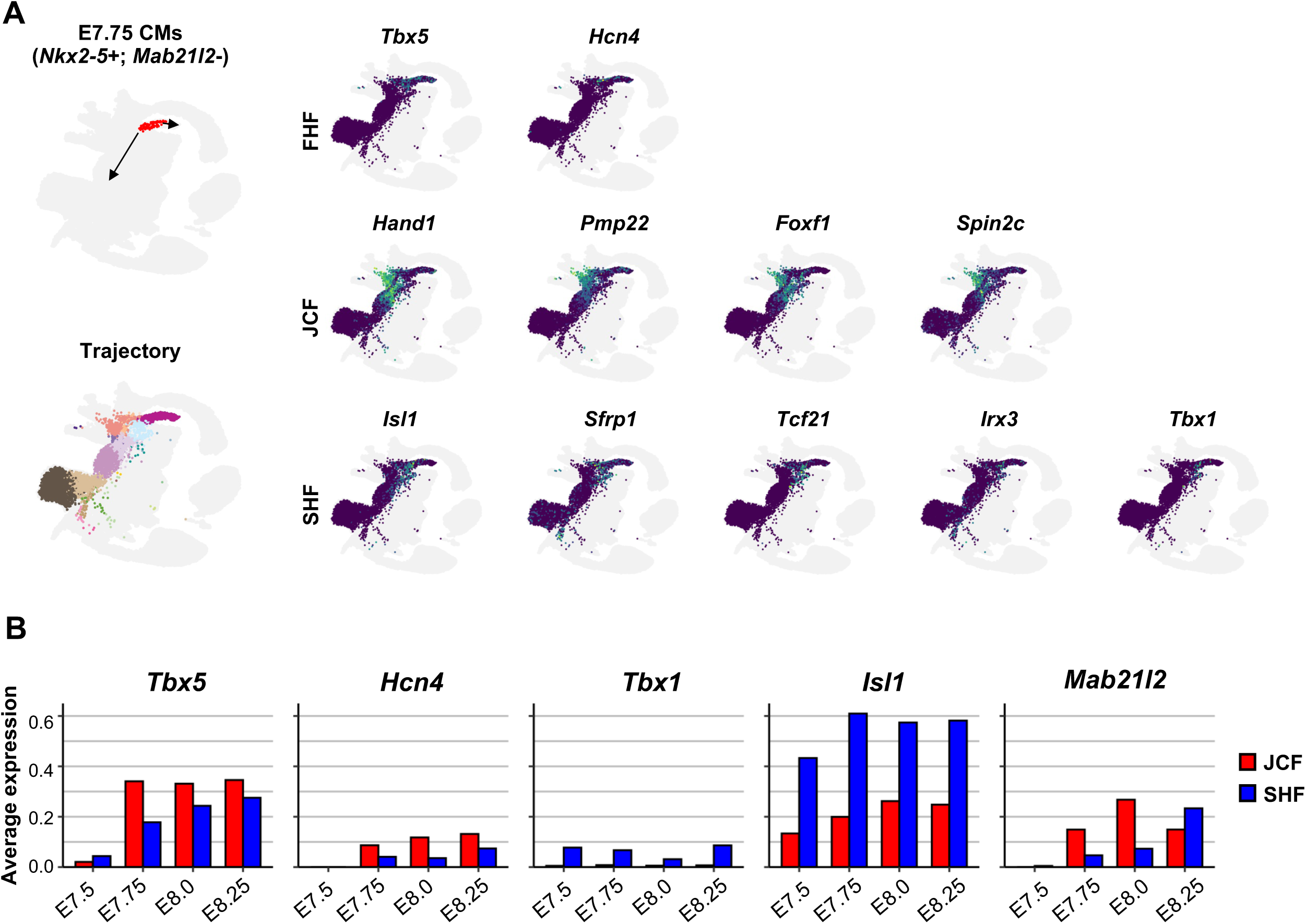
JCF and SHF progenitors contribute to the FHF **(A)** WOT analysis for early FHF progenitor population. UMAP layout from Pijuan-Sala et al. [12] is highlighted by cells belonging to the WOT predicted developmental trajectories for CM (E7.75 *Nkx2-5*+; *Mab21l2*- CM cells). Expression levels of representative marker genes for FHF, SHF JCF and SHF, projected onto the same UMAP. **(B)** Barplots indicates variable contribution of JCF and SHF to FHF. Bar hights represent the expression levels of FHF (*Tbx5, Hcn4*), SHF (*Tbx1, Isl1*) and JCF (*Mab21l2*) marker genes, averaged among single cells of each lineage.

**Figure S3:**
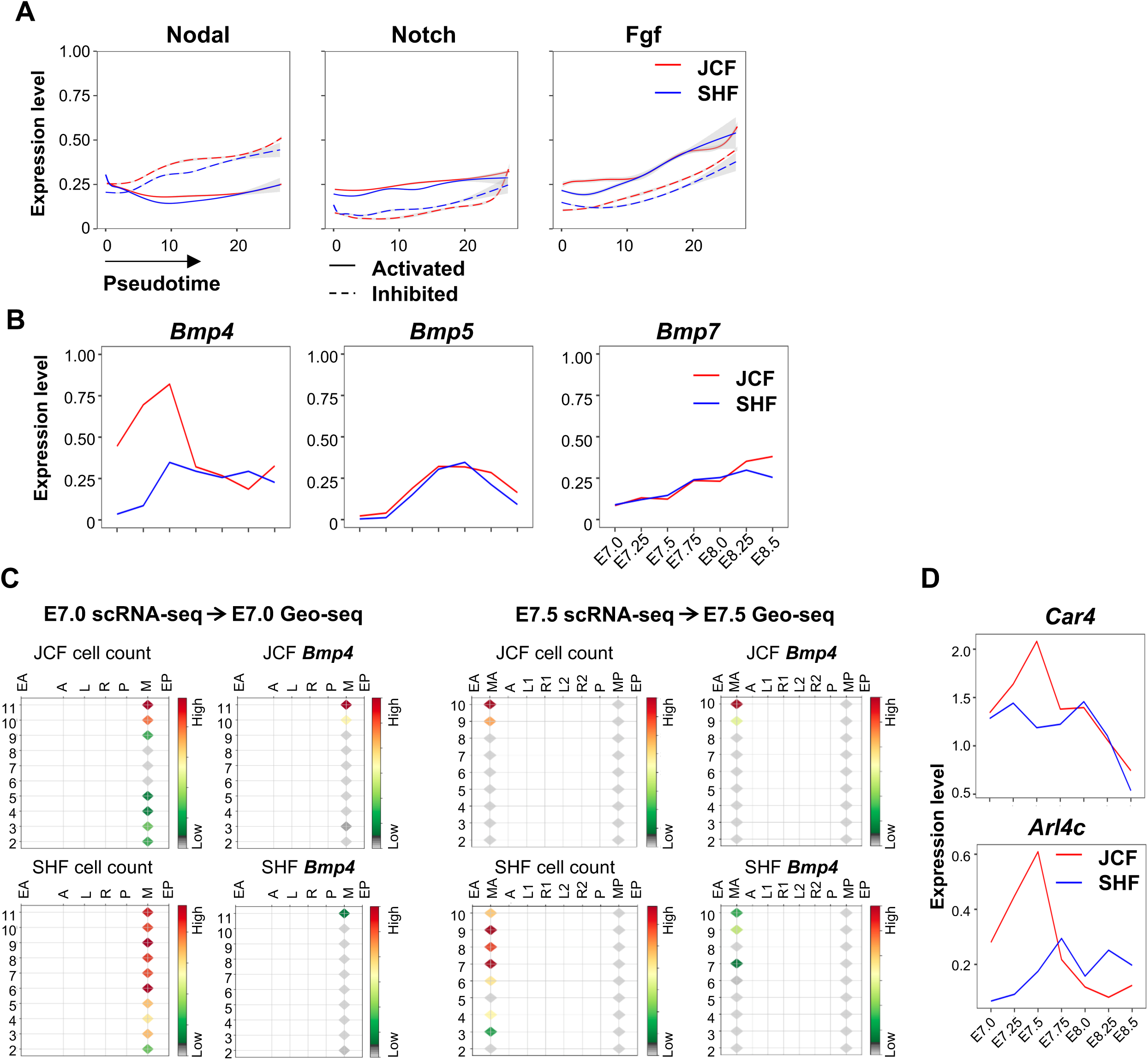
Differential gene expression and signaling activaity in JCF and SHF **(A)** Smoothened fitting curves showing expression levels of activated (solid line) and inhibited (dotted line) signaling target genes in JCF (red) and SHF (blue). **(B)** Dynamic expression of *Bmp4*/*5*/*7* in JCF and SHF between E7.0-E8.5 stages. **(C)** Corn plots showing spatial JCF (upper) and SHF single cell mapping at the E7.0 (left) and E7.5 (right) stages. Columns representing micro-dissected locations in germ layers and rows representing distal (slice 2) to proximal ends (slices 10/11) of the mouse embryos. Colors indicating the number of single cells mapped to each location, or the aggregated expression level of *Bmp4*. EA, anterior endoderm; EP, posterior endoderm; M, whole mesoderm; MA, anterior mesoderm; MP, posterior mesoderm; A, anterior ectoderm; P, posterior ectoderm; L, left lateral ectoderm; R, right lateral ectoderm. **(D)** Dynamic expression of the Bmp signaling target genes *Car4* and *Arl4c* in JCF and SHF between E7.0-E8.5 stages.

**Figure S4:**
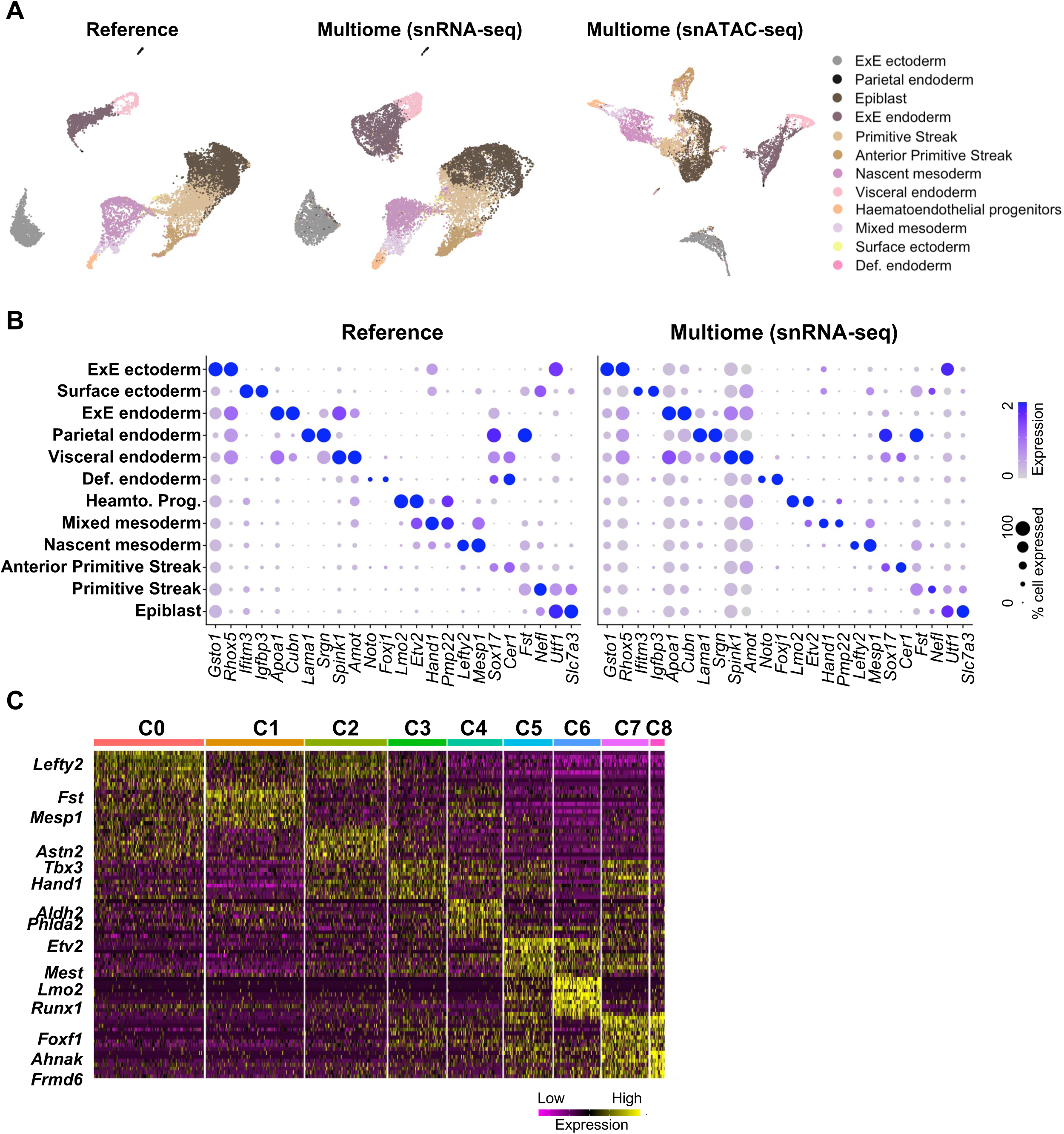
Data integration of E7.0 multi-omics snRNA-seq with reference scRNA-seq **(A)** UMAP layout of E7.0 snRNA-seq or snATAC-seq data. UMAPs of snRNA-seq was generated by integrating published [12] and multi-omics data of this study. Cells are colored by cell types. **(B)** Dot plot showing marker gene expression in reference and multi-omics dataset. **(C)** Heatmap showing expression levels of marker genes across mesodermal clusters of E7.0 mouse embryos.

**Figure S5:**
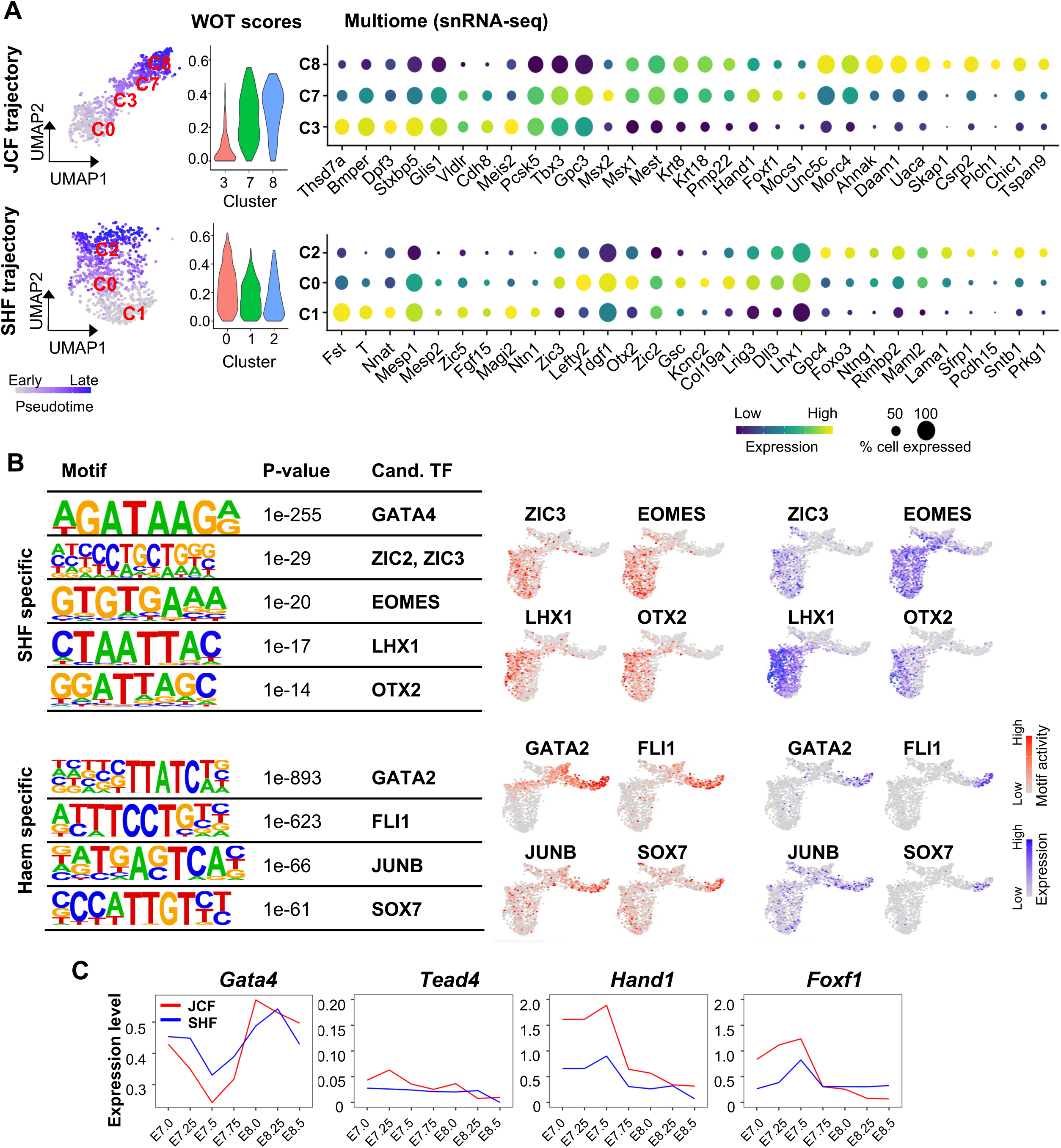
Identification of the key lineage specific TFs **(A)** UMAP layout (left) of the JCF (C0/3/7/8) and SHF (C1/0/2) developmental trajectories. Color shades indicating pseudotime for developmental stages. Violin plots (middle) showing the WOT score of C3/7/8 and C0/1/2 cells belong to JCF and SHF lineages, respectively. Dot plots (right) showing the marker gene expression for JCF specific clusters (C3/7/8) and SHF specific clusters (C1/0/2) in snRNA-seq. **(B)** Identification of top SHF (upper)/Haem (lower) specific DNA-binding motifs and corresponding candidate TFs. The SHF/Haem specific DAEs were defined by comparing C2/C6 with C1 snATAC-seq data using SnapATAC [30] ‘findDAR’ function. Motif calling was performed by the HOMER [31] ‘findMotifsGenome.pl’ function. Motif activity (colored in red) and TF expression (colored in blue) levels of trajectory specific candidate TFs, are overlaid on the UMAP layout from Figure 2A. **(C)** Dynamic expression of *Gata4*, *Tead4*, *Hand1* and *Foxf1* in JCF and SHF between E7.0-E8.5 developmental stages.

**Figure S6:**
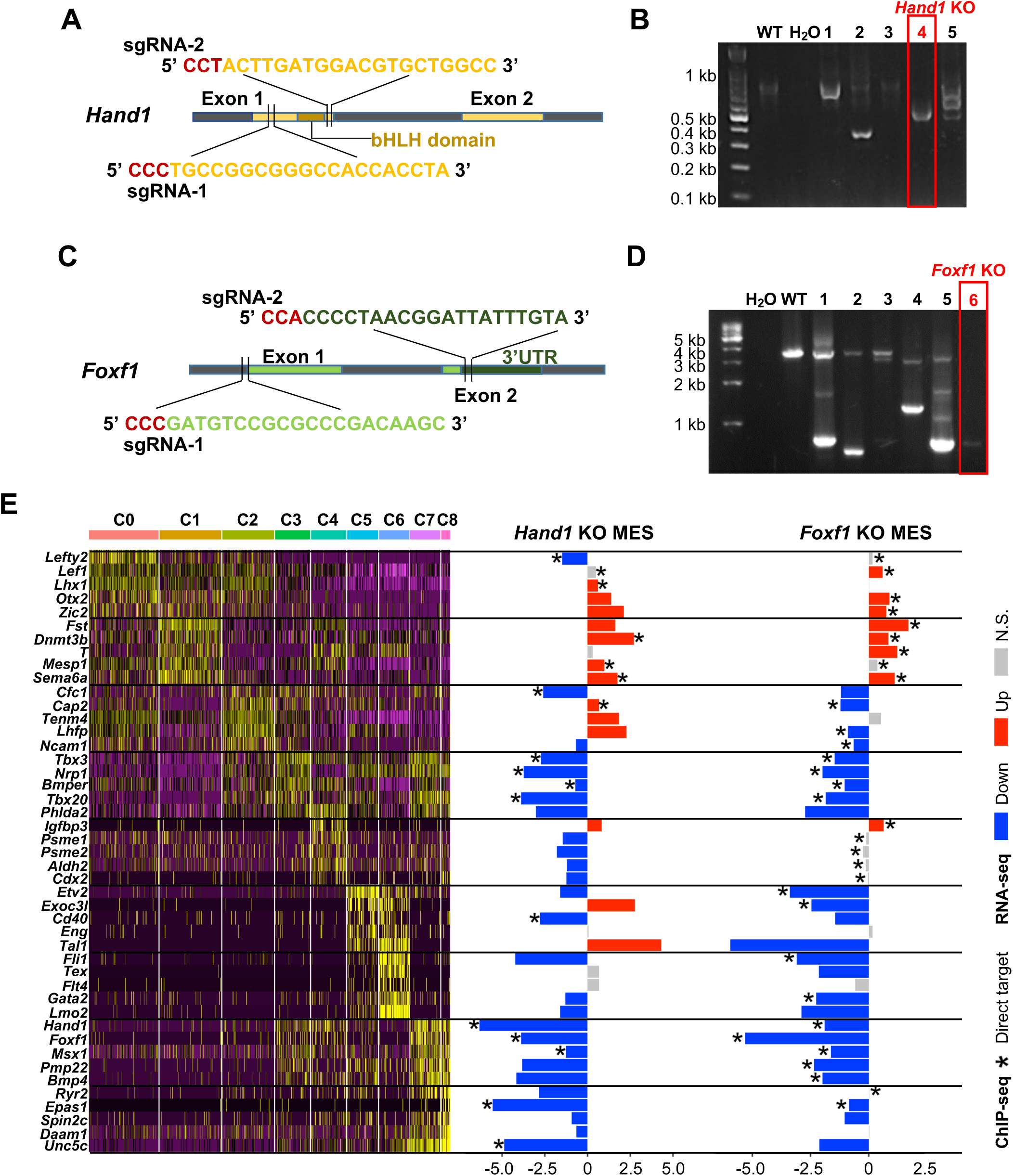
Generation of the Hand1 and Foxf1 KO ESC lines **(A,C)** Schematic diagram of the positions of sgRNAs targeting the CDS of *Hand1* (A) and *Foxf1* (C). **(B,D)** Genomic PCR analyses using the primer sets flanking the cleavage sites verifying the genomic DNA deletion of *Hand1* (B) and *Foxf1* (D). **(E)** Heatmap showing expression levels of the marker genes across mesodermal clusters of E7.0 mouse embryos (left). Bar plots showing expression FC upon *Hand1* and *Foxf1* KO (right). Asterisks indicating direct binding by the corresponding TFs.

**Figure S7:**
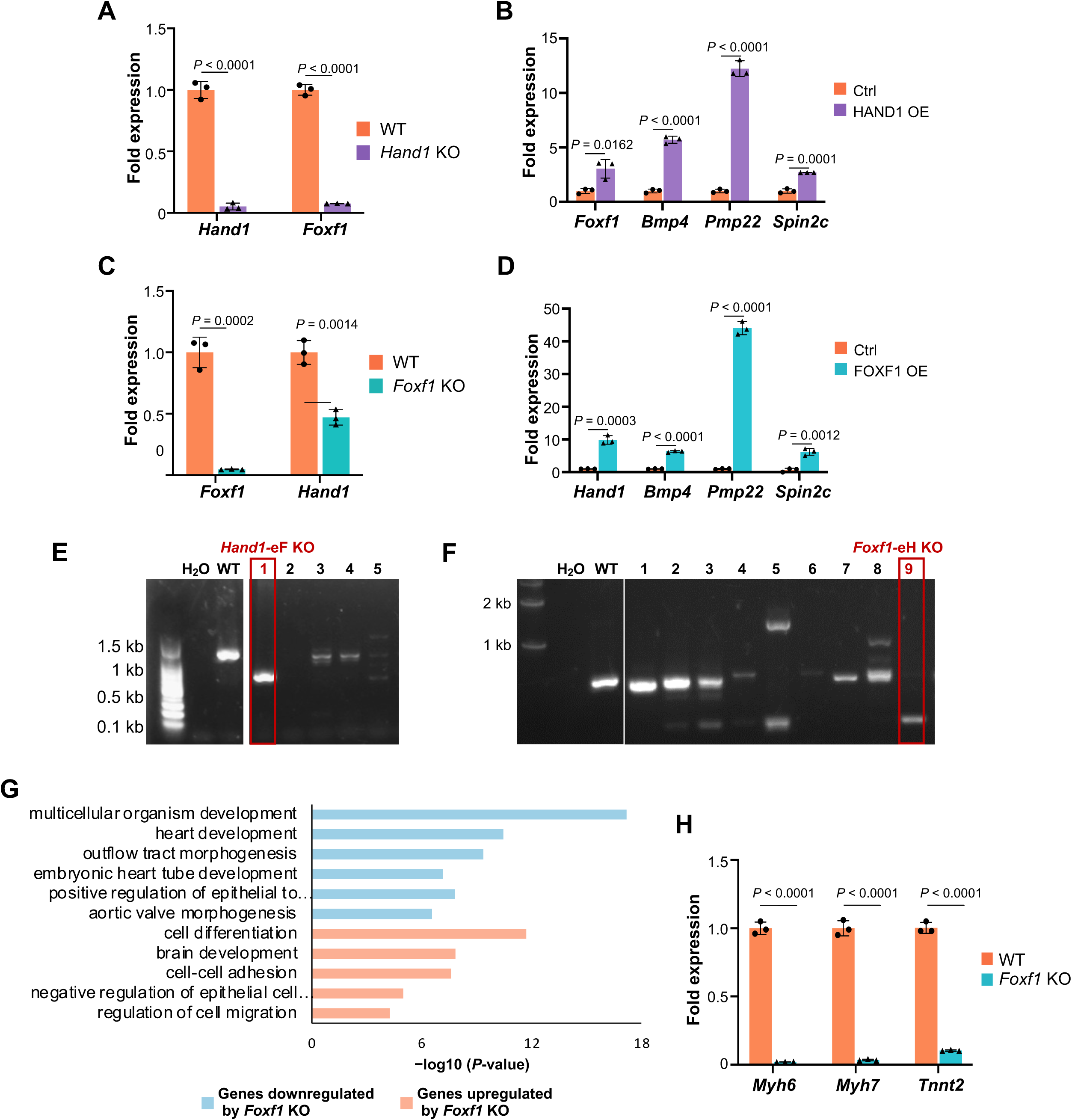
Mutual regulation between HAND1 and FOXF1 **(A-D)** RT-qPCR showing the levels of HAND1 and FOXF1 in *Hand1* and *Foxf1* KO, the EEM marker genes (*Foxf1, Bmp4, Pmp22* and *Spin2c*) after HAND1 and FOXF1 OE. **(E-F)** Genomic PCR analyses using the primer sets flanking the cleavage sites verifying the genomic DNA deletion of the *Hand1* enhancer (a) and the *Foxf1* enhancer (c). **(G)** Enriched GO terms of the genes up- or down-regulated after *Foxf1* KO. One-sided Fisher’s Exact test with Benjamini-Hochberg multiple testing correction was performed. **(H)** RT-qPCR showing that the expression levels of *Myh6*, *Myh7* and *Tnnt2* at CM stage are impaired by *Foxf1* KO. Data are the mean ± standard error of the mean from three independent experiments. Two-tailed unpaired Student’s *t*-test was performed.

**Figure S8:**
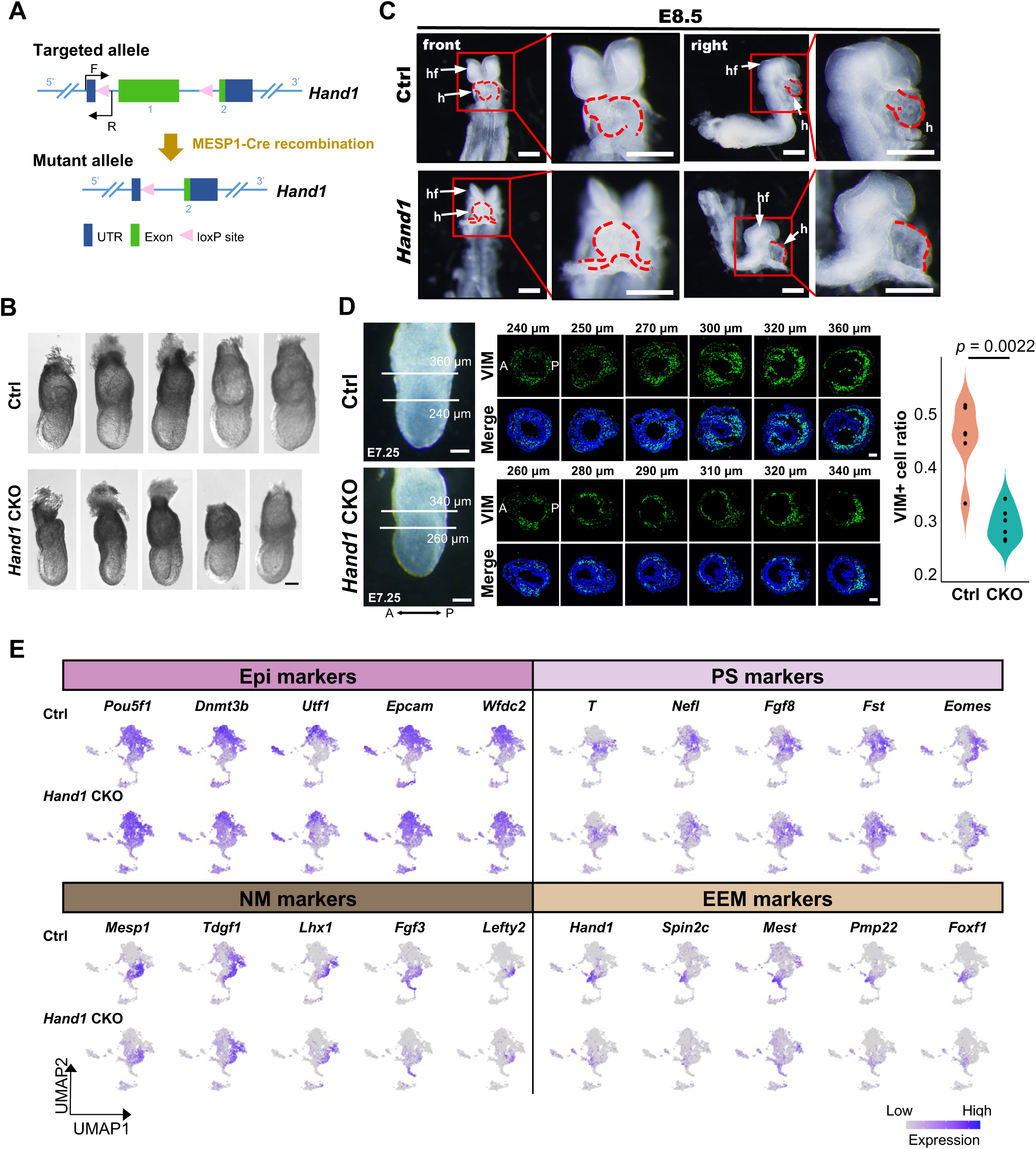
HAND1 is required for the expression of the EEM specific genes in vivo **(A)** Schematic diagram showing strategy for the mesodermal cell specific *Hand1* KO embryo generation. **(B)** Bright field images of representative E7.0 Ctrl and *Hand1* CKO embryos. Scale bar: 0.1 mm. **(C)** The bright field images of E8.5 Ctrl and *Hand1* CKO mouse embryos. The arrows indicating the embryonic heart (h) and head folds (hf). Scale bar: 500 μm. **(D)** The bright field image of E7.25 Ctrl and *Hand1* CKO mouse embryos (left), scale bar: 100 μm. Immunofluorescence staining of VIM (EEM marker) on serial embryo sections (middle), scale bar: 50 μm. Quantification of the proportion of VIM⁺ EEM cells, *P*-value was calculated using one-sided Mann-Whitney U test. **(E)** UMAP layout as in Figure 6C. Colors indicating expression levels of the marker genes of Epi, PS, NM and EEM.

**Figure S9:**
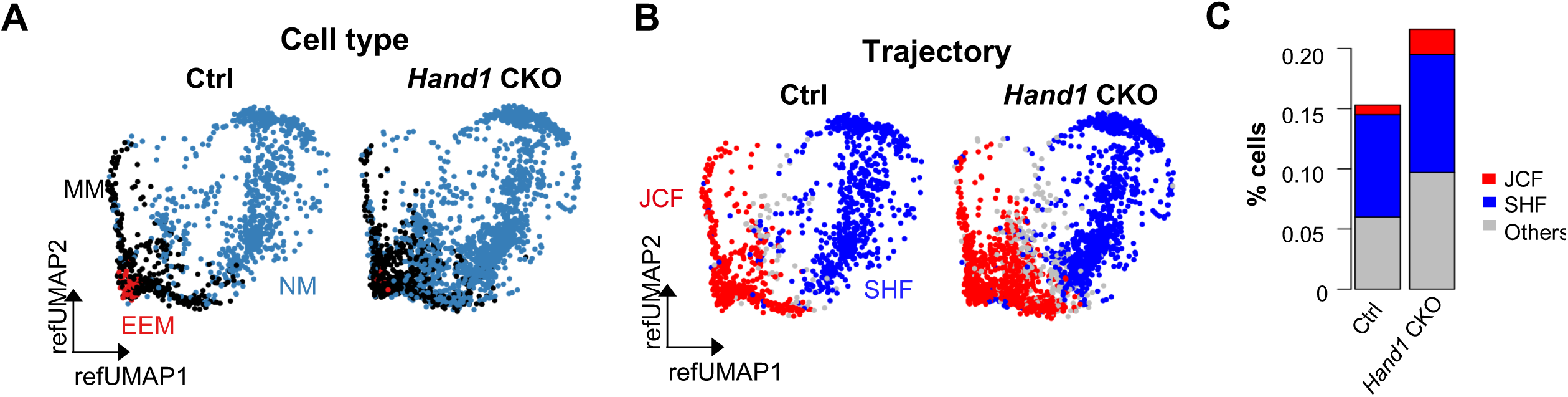
Integration of E7.0 progenitors and E7.0 Ctrl/Hand1 KO scRNA-seq **(A-B)** Projection of E7.0 scRNA-seq onto the reference UMAP structure from Figure 1D. Reference UMAP layout for E7.0 CM trajectory cells colored by cell type (A) and trajectory (B). **(C)** Bar plot showing the percentage of the JCF and SHF trajectories in NM of Ctrl and *Hand1* CKO mice.

## Supplemental Tables

**Table S1.1:**
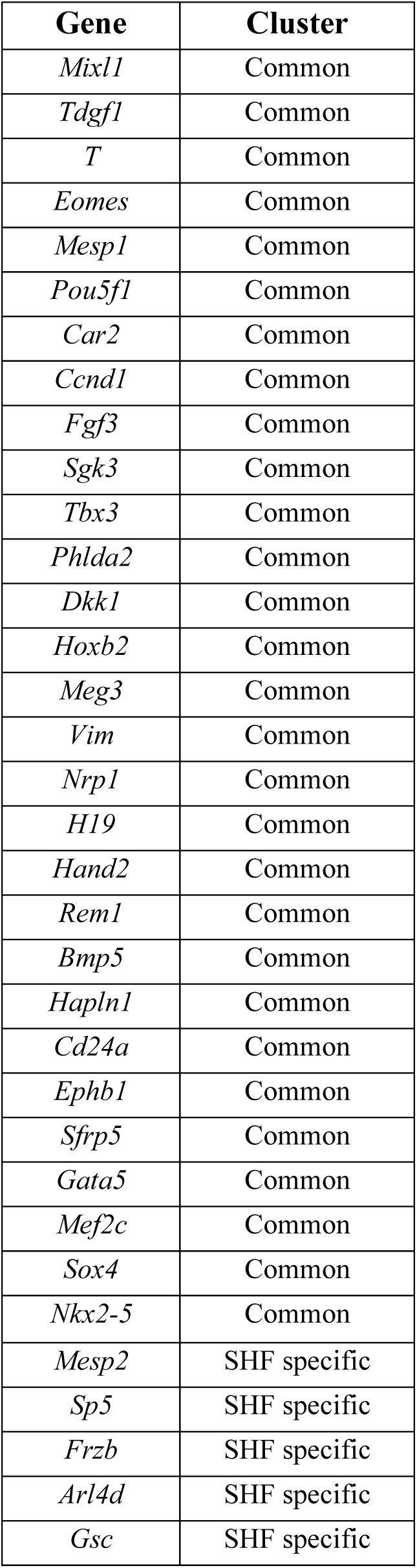
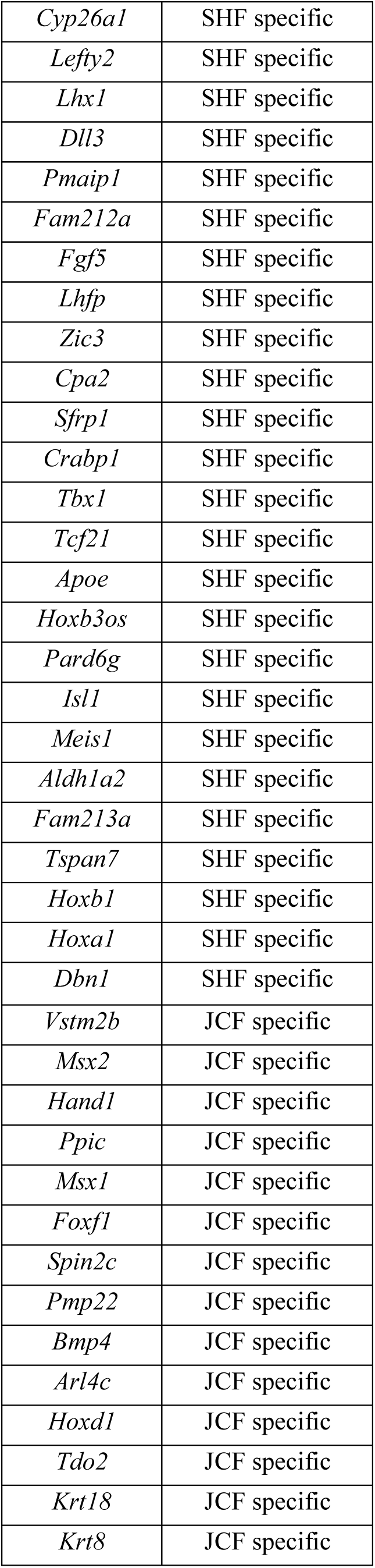
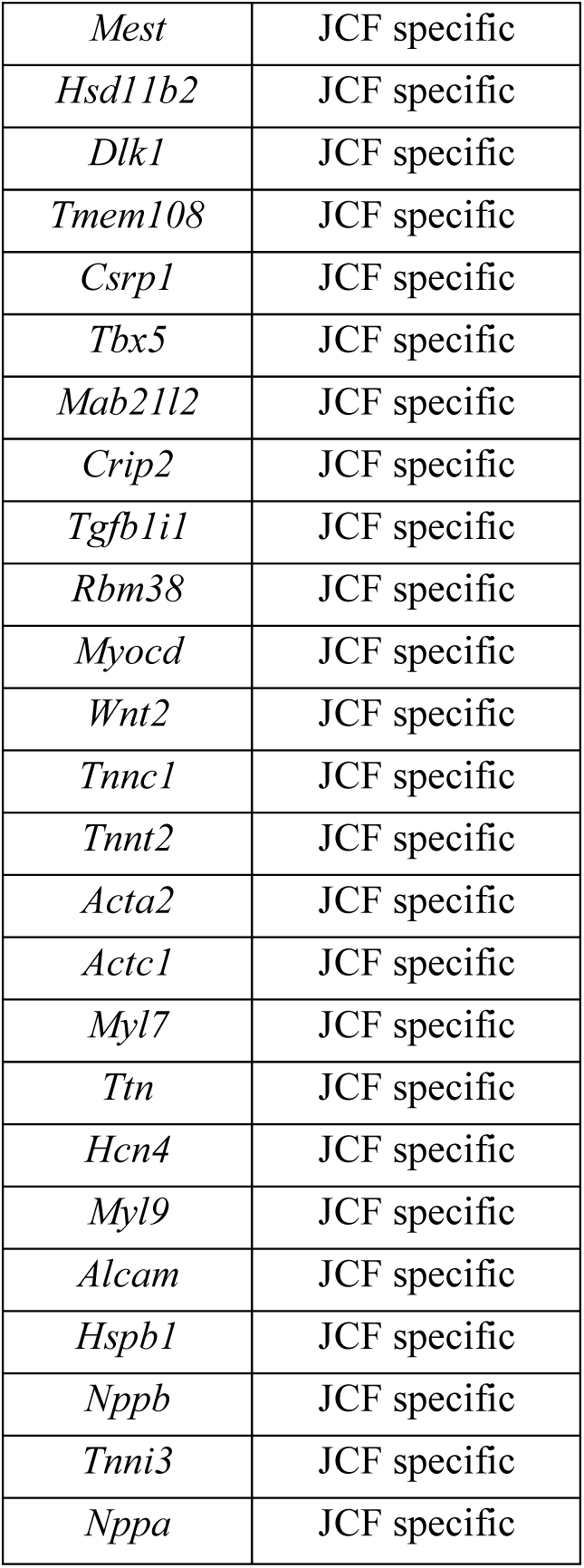
Pseudotime-dependent genes for JCF/SHF trajectoyies. Related to. **Figure 1F**

**Table S1.2:**
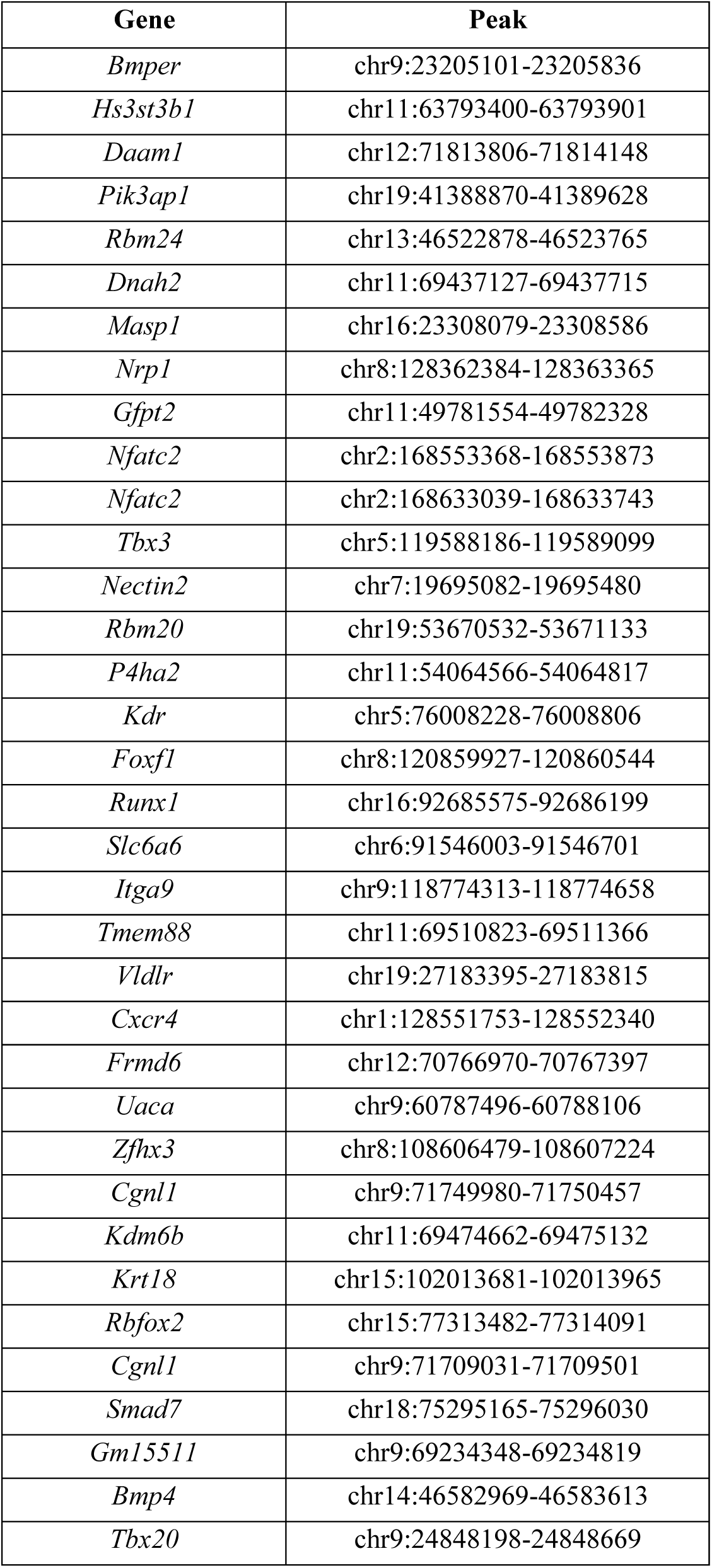
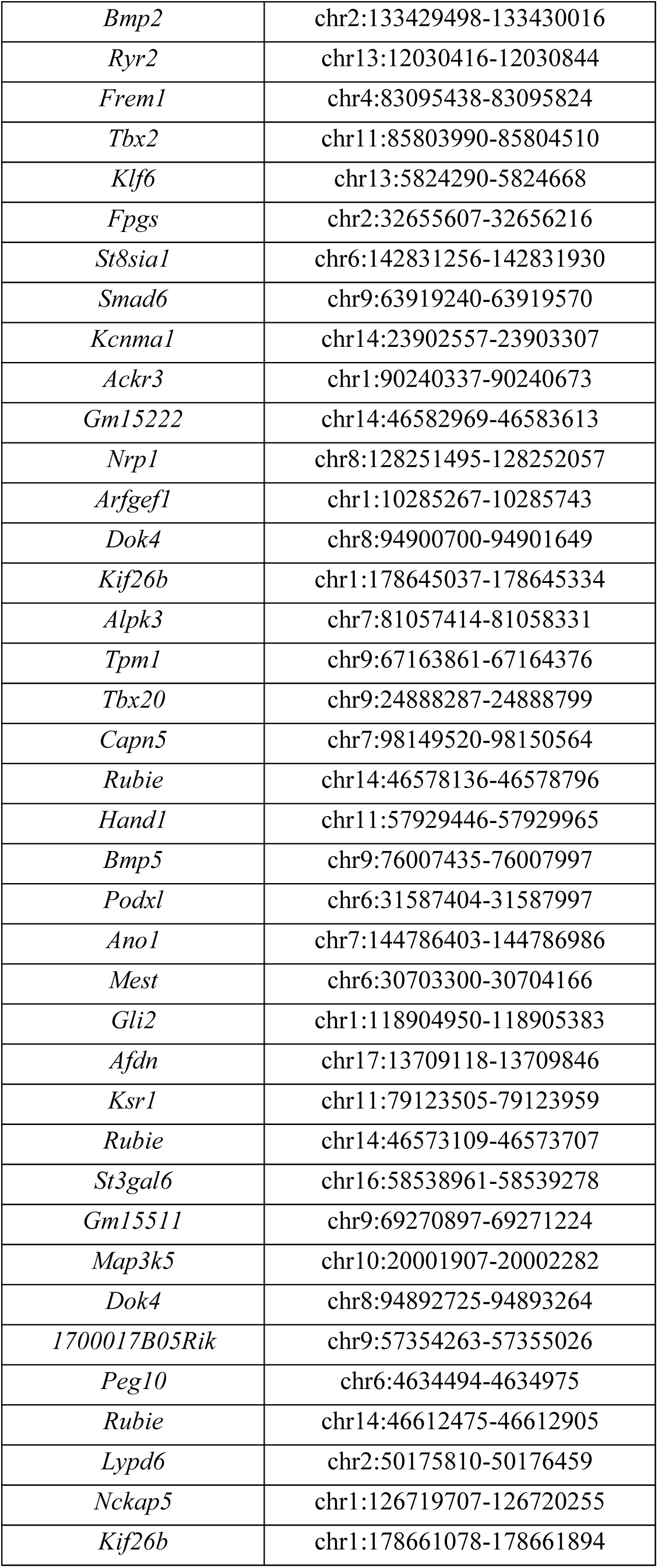
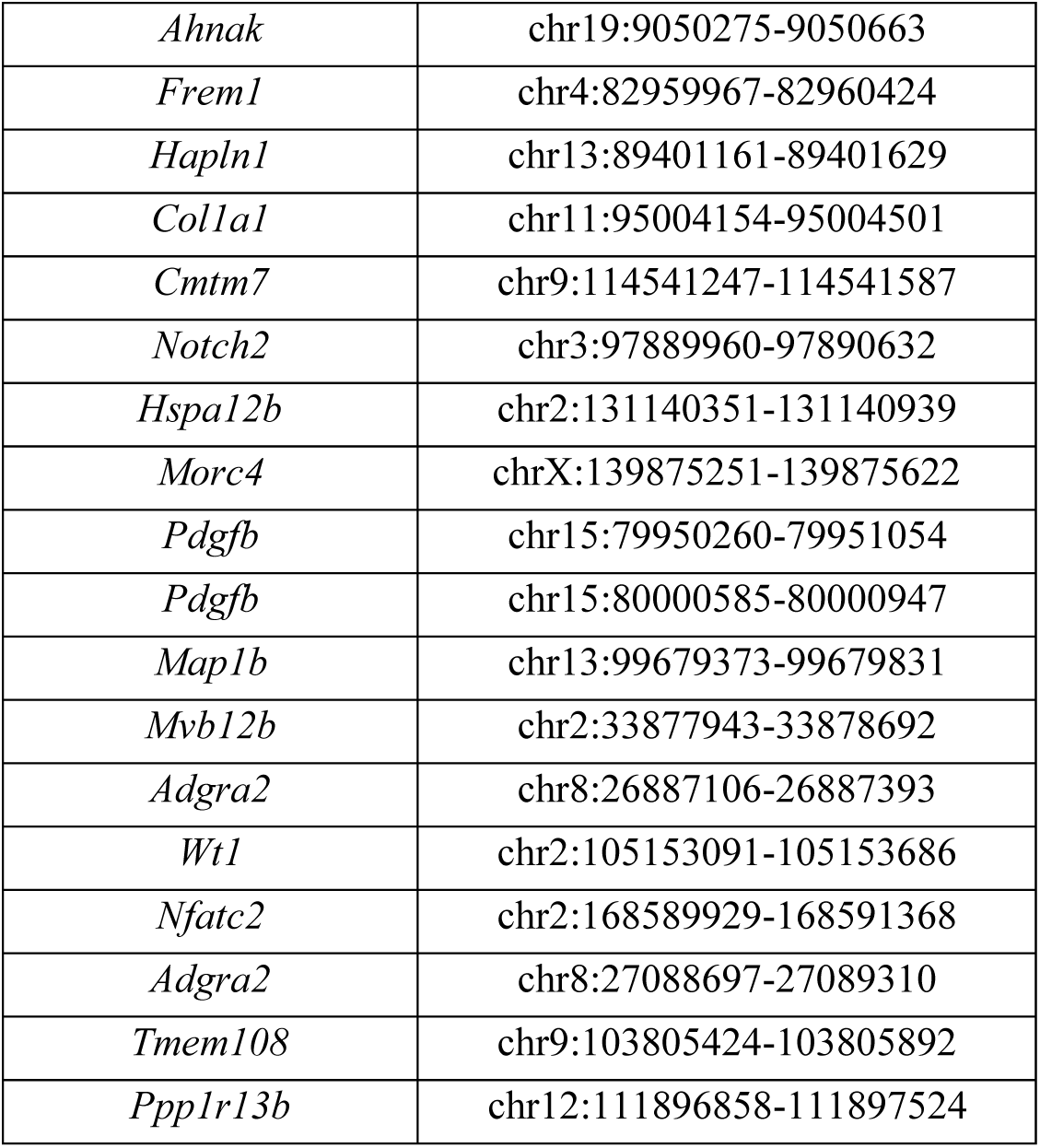
Gene-enhancer pairs in E7.0 mesoderm pseudotime trajectories. Related to. **Figure 2G**

Table S2. ***Hand1*- and *Foxf1*-KO affected genes in MES cells. Attached as an excel file.**

**Table S3:**
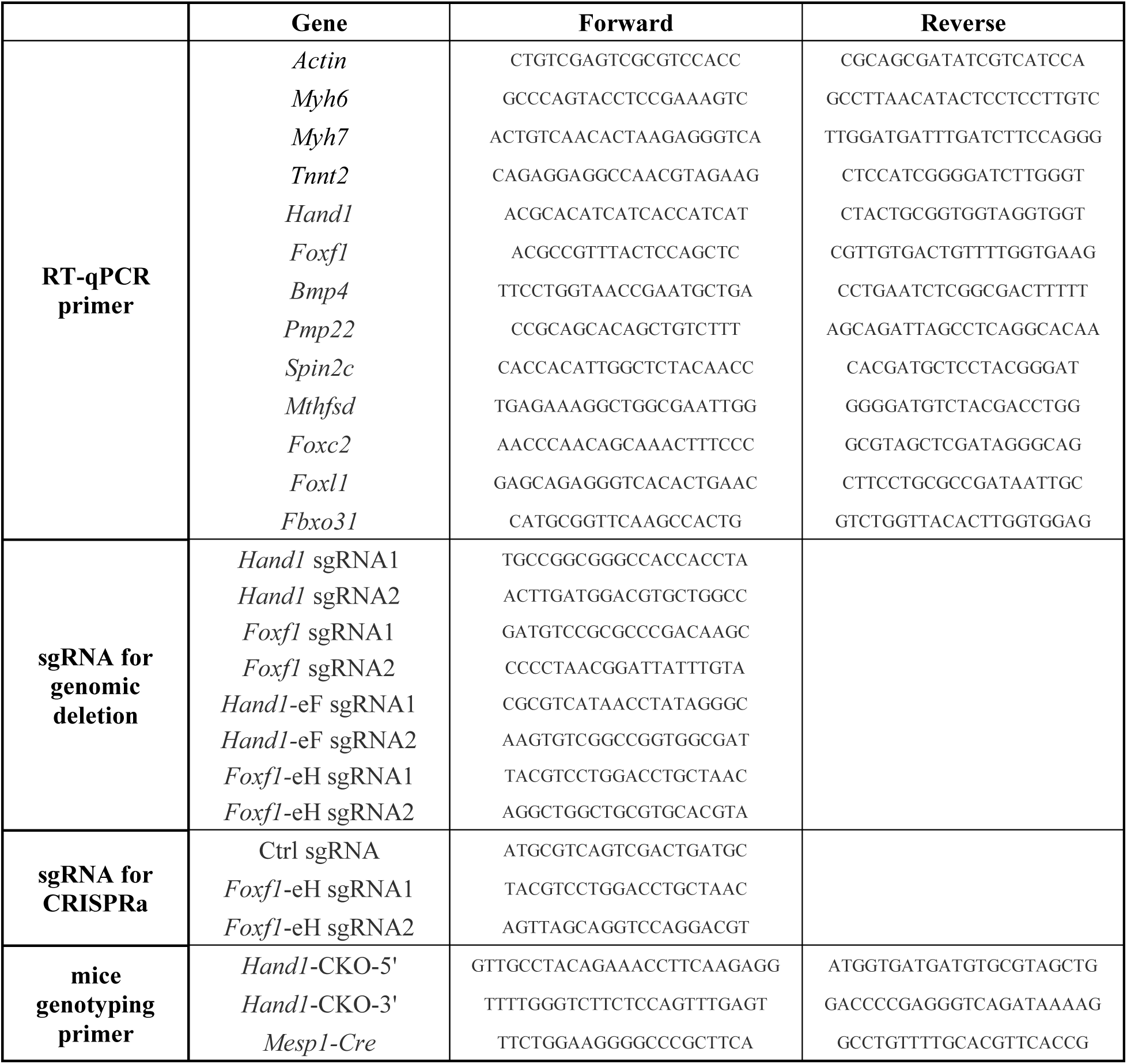
Primers/oligos used in the current study.

## Notes

### Competing Interest Statement

The authors have declared no competing interest.

### Summary of Updates

We provided additional computational and experimental data to strengthen the analyses about cardiac lineage trajectories and TF functions.

## Reference

[1] M. Buckingham, S. Meilhac, S. Zaffran, Nat Rev Genet 2005, 6 (11), 826, 10.1038/nrg1710.

[2] R. G. Kelly, Dev Cell 2023, 58 (4), 257, 10.1016/j.devcel.2023.01.010.

[3] R. Abu-Issa, M. L. Kirby, Annu Rev Cell Dev Biol 2007, 23, 45, 10.1146/annurev.cellbio.23.090506.123331; M. Parameswaran, P. P. Tam, *Dev Genet* 1995, *17* (1), 16, 10.1002/dvg.1020170104.

4. R. C. V. Tyser, X. Ibarra-Soria, K. McDole, S. Arcot Jayaram, J. Godwin, T. A. H. van den Brand, A. M. A. Miranda, A. Scialdone, P. J. Keller, J. C. Marioni, S. Srinivas, Science 2021, *371* (6533), 10.1126/science.abb2986.

[5] C. L. Cai, X. Liang, Y. Shi, P. H. Chu, S. L. Pfaff, J. Chen, S. Evans, Dev Cell 2003, 5 (6), 877, 10.1016/s1534-5807(03)00363-0.

[6] Q. Zhang, D. Carlin, F. Zhu, P. Cattaneo, T. Ideker, S. M. Evans, J. Bloomekatz, N. C. Chi, Circ Res 2021, 129 (4), 474, 10.1161/CIRCRESAHA.121.318943.

[7] Y. Saga, S. Miyagawa-Tomita, A. Takagi, S. Kitajima, J. Miyazaki, T. Inoue, Development 1999, 126 (15), 3437, 10.1242/dev.126.15.3437.

[8] W. P. Devine, J. D. Wythe, M. George, K. Koshiba-Takeuchi, B. G. Bruneau, Elife 2014, 3, 10.7554/eLife.03848; F. Lescroart, S. Chabab, X. Lin, S. Rulands, C. Paulissen, A. Rodolosse, H. Auer, Y. Achouri, C. Dubois, A. Bondue, B. D. Simons, C. Blanpain, *Nat Cell Biol* 2014, *16* (9), 829, 10.1038/ncb3024.

[9] F. Lescroart, X. Wang, X. Lin, B. Swedlund, S. Gargouri, A. Sanchez-Danes, V. Moignard, C. Dubois, C. Paulissen, S. Kinston, B. Gottgens, C. Blanpain, Science 2018, 359 (6380), 1177, 10.1126/science.aao4174.

[10] K. Ivanovitch, P. Soro-Barrio, P. Chakravarty, R. A. Jones, D. M. Bell, S. N. Mousavy Gharavy, D. Stamataki, J. Delile, J. C. Smith, J. Briscoe, PLoS Biol 2021, 19 (5), e3001200, 10.1371/journal.pbio.3001200.

[11] G. Schiebinger, J. Shu, M. Tabaka, B. Cleary, V. Subramanian, A. Solomon, J. Gould, S. Liu, S. Lin, P. Berube, L. Lee, J. Chen, J. Brumbaugh, P. Rigollet, K. Hochedlinger, R. Jaenisch, A. Regev, E. S. Lander, Cell 2019, 176 (4), 928, 10.1016/j.cell.2019.01.006.

[12] B. Pijuan-Sala, J. A. Griffiths, C. Guibentif, T. W. Hiscock, W. Jawaid, F. J. Calero-Nieto, C. Mulas, X. Ibarra-Soria, R. C. V. Tyser, D. L. L. Ho, W. Reik, S. Srinivas, B. D. Simons, J. Nichols, J. C. Marioni, B. Gottgens, Nature 2019, 566 (7745), 490, 10.1038/s41586-019-0933-9.

[13] M. H. Dominguez, A. L. Krup, J. M. Muncie, B. G. Bruneau, Cell 2023, 186 (3), 479, 10.1016/j.cell.2023.01.001.

[14] X. Yang, B. Hu, J. Liao, Y. Qiao, Y. Chen, Y. Qian, S. Feng, F. Yu, J. Dong, Y. Hou, H. Xu, R. Wang, G. Peng, J. Li, F. Tang, N. Jing, Cell Res 2019, 29 (11), 911, 10.1038/s41422-019-0234-8.

[15] J. A. Wamstad, J. M. Alexander, R. M. Truty, A. Shrikumar, F. Li, K. E. Eilertson, H. Ding, J. N. Wylie, A. R. Pico, J. A. Capra, G. Erwin, S. J. Kattman, G. M. Keller, D. Srivastava, S. S. Levine, K. S. Pollard, A. K. Holloway, L. A. Boyer, B. G. Bruneau, Cell 2012, 151 (1), 206, 10.1016/j.cell.2012.07.035.

[16] A. He, F. Gu, Y. Hu, Q. Ma, L. Y. Ye, J. A. Akiyama, A. Visel, L. A. Pennacchio, W. T. Pu, Nat Commun 2014, 5, 4907, 10.1038/ncomms5907.

[17] X. Lin, B. Swedlund, M. N. Ton, S. Ghazanfar, C. Guibentif, C. Paulissen, E. Baudelet, E. Plaindoux, Y. Achouri, E. Calonne, C. Dubois, W. Mansfield, S. Zaffran, J. C. Marioni, F. Fuks, B. Gottgens, F. Lescroart, C. Blanpain, Nat Cell Biol 2022, 24 (7), 1114, 10.1038/s41556-022-00947-3.

[18] A. B. Firulli, D. G. McFadden, Q. Lin, D. Srivastava, E. N. Olson, Nat Genet 1998, 18 (3), 266, 10.1038/ng0398-266.

[19] P. Riley, L. Anson-Cartwright, J. C. Cross, Nat Genet 1998, 18 (3), 271, 10.1038/ng0398-271.

[20] J. A. Farrell, Y. Wang, S. J. Riesenfeld, K. Shekhar, A. Regev, A. F. Schier, Science 2018, 360 (6392), 10.1126/science.aar3131.

[21] M. Mittnenzweig, Y. Mayshar, S. Cheng, R. Ben-Yair, R. Hadas, Y. Rais, E. Chomsky, N. Reines, A. Uzonyi, L. Lumerman, A. Lifshitz, Z. Mukamel, A. H. Orenbuch, A. Tanay, Y. Stelzer, Cell 2021, 184 (11), 2825, 10.1016/j.cell.2021.04.004; C. Qiu, J. Cao, B. K. Martin, T. Li, I. C. Welsh, S. Srivatsan, X. Huang, D. Calderon, W. S. Noble, C. M. Disteche, S. A. Murray, M. Spielmann, C. B. Moens, C. Trapnell, J. Shendure, *Nat Genet* 2022, *54* (3), 328, 10.1038/s41588-022-01018-x.

[22] T. Y. de Soysa, S. S. Ranade, S. Okawa, S. Ravichandran, Y. Huang, H. T. Salunga, A. Schricker, A. Del Sol, C. A. Gifford, D. Srivastava, Nature 2019, 572 (7767), 120, 10.1038/s41586-019-1414-x.

[23] S. M. Meilhac, M. Esner, R. G. Kelly, J. F. Nicolas, M. E. Buckingham, Dev Cell 2004, 6 (5), 685, 10.1016/s1534-5807(04)00133-9.

[24] M. Zheng, S. Erhardt, D. Ai, J. Wang, Int J Mol Sci 2021, 22 (18), 10.3390/ijms22189835.

[25] M. Fleury, A. Eliades, P. Carlsson, G. Lacaud, V. Kouskoff, Development 2015, 142 (19), 3307, 10.1242/dev.124685.

[26] Z. Xue, K. Huang, C. Cai, L. Cai, C. Y. Jiang, Y. Feng, Z. Liu, Q. Zeng, L. Cheng, Y. E. Sun, J. Y. Liu, S. Horvath, G. Fan, Nature 2013, 500 (7464), 593, 10.1038/nature12364.

[27] C. A. Risebro, N. Smart, L. Dupays, R. Breckenridge, T. J. Mohun, P. R. Riley, Development 2006, 133 (22), 4595, 10.1242/dev.02625.

[28] H. Guo, C. Hang, B. Lin, Z. Lin, H. Xiong, M. Zhang, R. Lu, J. Liu, D. Shi, D. Xie, Y. Liu, D. Liang, J. Yang, Y. H. Chen, Stem Cell Res Ther 2024, 15 (1), 31, 10.1186/s13287-024-03649-9.

[29] A. Sampath Kumar, L. Tian, A. Bolondi, A. A. Hernandez, R. Stickels, H. Kretzmer, E. Murray, L. Wittler, M. Walther, G. Barakat, L. Haut, Y. Elkabetz, E. Z. Macosko, L. Guignard, F. Chen, A. Meissner, Nat Genet 2023, 55 (7), 1176, 10.1038/s41588-023-01435-6.

[30] R. Fang, S. Preissl, Y. Li, X. Hou, J. Lucero, X. Wang, A. Motamedi, A. K. Shiau, X. Zhou, F. Xie, E. A. Mukamel, K. Zhang, Y. Zhang, M. M. Behrens, J. R. Ecker, B. Ren, Nat Commun 2021, 12 (1), 1337, 10.1038/s41467-021-21583-9.

[31] S. Heinz, C. Benner, N. Spann, E. Bertolino, Y. C. Lin, P. Laslo, J. X. Cheng, C. Murre, H. Singh, C. K. Glass, Mol Cell 2010, 38 (4), 576, 10.1016/j.molcel.2010.05.004.

[32] J. Nie, M. Jiang, X. Zhang, H. Tang, H. Jin, X. Huang, B. Yuan, C. Zhang, J. C. Lai, Y. Nagamine, D. Pan, W. Wang, Z. Yang, Cell Rep 2015, 13 (4), 723, 10.1016/j.celrep.2015.09.043.

[33] Y. Hao, S. Hao, E. Andersen-Nissen, W. M. Mauck, 3rd, S. Zheng, A. Butler, M. J. Lee, A. J. Wilk, C. Darby, M. Zager, P. Hoffman, M. Stoeckius, E. Papalexi, E. P. Mimitou, J. Jain, A. Srivastava, T. Stuart, L. M. Fleming, B. Yeung, A. J. Rogers, J. M. McElrath, C. A. Blish, R. Gottardo, P. Smibert, R. Satija, Cell 2021, 184 (13), 3573, 10.1016/j.cell.2021.04.048.

[34] C. S. McGinnis, L. M. Murrow, Z. J. Gartner, Cell Syst 2019, 8 (4), 329, 10.1016/j.cels.2019.03.003.

[35] B. Langmead, S. L. Salzberg, Nat Methods 2012, 9 (4), 357, 10.1038/nmeth.1923.

[36] F. Ramirez, D. P. Ryan, B. Gruning, V. Bhardwaj, F. Kilpert, A. S. Richter, S. Heyne, F. Dundar, T. Manke, Nucleic Acids Res 2016, 44 (W1), W160, 10.1093/nar/gkw257.

[37] Y. Zhang, T. Liu, C. A. Meyer, J. Eeckhoute, D. S. Johnson, B. E. Bernstein, C. Nusbaum, R. M. Myers, M. Brown, W. Li, X. S. Liu, Genome Biol 2008, 9 (9), R137, 10.1186/gb-2008-9-9-r137.

[38] G. Peng, S. Suo, G. Cui, F. Yu, R. Wang, J. Chen, S. Chen, Z. Liu, G. Chen, Y. Qian, P. P. L. Tam, J. J. Han, N. Jing, Nature 2019, 572 (7770), 528, 10.1038/s41586-019-1469-8.

[39] S. J. Pan, I. W. Tsang, J. T. Kwok, Q. Yang, IEEE Trans Neural Netw 2011, 22 (2), 199, 10.1109/TNN.2010.2091281.

[40] C. Trapnell, D. Cacchiarelli, J. Grimsby, P. Pokharel, S. Li, M. Morse, N. J. Lennon, K. J. Livak, T. S. Mikkelsen, J. L. Rinn, Nat Biotechnol 2014, 32 (4), 381, 10.1038/nbt.2859.

[41] T. Wu, E. Hu, S. Xu, M. Chen, P. Guo, Z. Dai, T. Feng, L. Zhou, W. Tang, L. Zhan, X. Fu, S. Liu, X. Bo, G. Yu, Innovation (Camb) 2021, 2 (3), 100141, 10.1016/j.xinn.2021.100141.

[42] A. N. Schep, B. Wu, J. D. Buenrostro, W. J. Greenleaf, Nat Methods 2017, 14 (10), 975, 10.1038/nmeth.4401.

[43] C. Stringer, T. Wang, M. Michaelos, M. Pachitariu, Nat Methods 2021, 18 (1), 100, 10.1038/s41592-020-01018-x.

[44] H. Peng, A. Bria, Z. Zhou, G. Iannello, F. Long, Nat Protoc 2014, 9 (1), 193, 10.1038/nprot.2014.011.

